# Common mycelial network modulates neighbour-primed plant defences against foliar pathogens by co-opting distinct inter-plant metabolic and biotic stress responses

**DOI:** 10.1101/2024.12.03.626652

**Authors:** Zigmunds Orlovskis, Ilva Nakurte, Ēriks Voroņins, Annija Kotova, Daniels Pugačevskis, Dawood Shah, Annija Andersone, Kārlis Trevors Blūms, Ivars Silamiķelis, Soon-Jae Lee

## Abstract

Arbuscular mycorrhizal fungi (AMF) are ubiquitous root symbionts that form common mycelial networks (CMN), linking multiple plants underground. CMN are hypothesized to play a role for information exchange between plants for neighbour-primed defences. However, the key transcriptomic and metabolome responses in receiver plant associated with inter-plant CMN connections remain yet to be elucidated. Additionally, the confounding effects of hyphal damage from CMN disconnection have not been clearly resolved. To uncover the contribution of CMN integrity to neighbour-primed plant defences, we used model AMF *Rhizophagus irregularis* to inter-connect two *Medicago truncatula* plants and explored the effect of sender wounding and *flg22* elicitation on receiver plants’ leaf responses and pathogen tolerance. For the first time, we demonstrate that changes in receiver’s biotic stress and defence signalling pathways rely on CMN-mediated inter-plant signals, not on mycelial network damage. This response was associated with distinct leaf isoprenoid production, including volatile monoterpenes and triterpene saponins. Furthermore, CMN-mediated signals from stressed senders enhanced receiver resistance to *Fusarium sporotrichoides* whilst simultaneously increasing susceptibility to *Botrytis cinerea*. Our findings highlight the critical role of CMN in inter-plant signalling for pathogen-specific susceptibility and resistance which can be a key for understanding plant community-level defence in nature and agroecosystems.

## Introduction

Exchanging danger signals between plants provides an effective strategy for priming the defences of receivers and preparing them for imminent pest or pathogen attacks that have already affected their neighbours. Such inter-plant communication may contribute to enhanced community-level resilience against biotic stress and presents a promising trait for future crop improvements and agricultural engineering. Upon perception of pest and pathogen elicitors such as herbivore eggs^1^ or bacterial virulence proteins^2^ plants are known to elevate pathogen resistance in their neighbours via airborne volatile or soil-mediated signals. Furthermore, ubiquitous plant root symbionts, such as arbuscular mycorrhizal fungi (AMF), are capable of inter-connecting root systems from multiple plant hosts, effectively forming underground common mycelial networks (CMN) which are hypothesized to function as information superhighway in transferring diverse signals from plant-to-plant in response to different sender plant stimuli^3,4^. Given that the vast majority of land plants form AMF symbiosis^5^, testing this hypothesis is paramount to understanding the ecological roles and mechanisms how CMN can contribute to the health of wild plants and crops.

While a few studies^6–8^ have implicated CMN for nutrient redistribution among the connected plants, functional studies on CMN-mediated inter-plant signals involved in plant defences and stress tolerance have been extremely scarce to date. Moreover, in all previous studies, which used CMN disruption^9,10^ to test the hypothesis whether CMN mediates stress-induced inter-plant signals, the effect from CMN disruption was confounded by mechanical damage of the symbiotic hyphal network. Therefore, the neighbour plant responses to signals associated with disconnection of inter-plant CMN were not decoupled from potential plant responses resulting from CMN hyphal damage. Another key shortcoming of the previous studies is the lack of a control for inducible sender or donor plant signal. For example, previous studies in tomato^9^ and pea^10^ did not include controls for pathogen non-infected or aphid non-infested sender plants, impeding non-biased conclusions about participation of CMN in stress-inducible as opposed to constitutive inter-plant signal exchange.

In the present study, we sought to overcome the limitations by previous studies and test two working hypotheses: whether inter-plant CMN connection, firstly, contributes to distinct receiver responses to stress-induced inter-plant signals and, secondly, mediates neighbour-primed defences against plant pathogens. To decouple the effect of CMN disruption from the symbiotic hyphal network damage, we set an experimental control where the disruption of mycorrhizal hyphal network occurs without perturbing the inter-plant CMN continuity. We used model AMF *Rhizophagus irregularis* to establish CMN between two *Medicago truncatula* plants, then subjected the leaves of signal senders to known plant defence elicitors with non-stimulated sender controls and analysed transcriptional and metabolic responses in naïve signal receiver plants. Intriguingly, we discovered CMN-connection dependent transcriptional changes in receivers, involving strong modulation of multiple plant biotic stress and defence signalling pathways. Moreover, leaf isoprenoid, especially monoterpene and triterpene saponin, production was elevated in receivers with the intact inter-plant CMN compared to the cut CMN independently from hyphal damage, strongly supporting our first hypothesis. Surprisingly, after the elicitation of sender plants, CMN-connected receiver plants exhibited heightened susceptibility to the fungal pathogen *Botrytis cinerea*, while simultaneously showing increased resistance to *Fusarium sporotrichoides*. This finding aligns with our second hypothesis whilst revealing previously undocumented species-specific effects in neighbour-modulated pathogen susceptibility within CMN-linked plant systems.

## Results

### Wounding combined with flg22 induces local leaf responses and systemic acquired resistance to Botrytis cinerea

To test the responses of signal receivers to sender stimuli, we first wished to identify stimuli that would induce leaf responses in the signal senders themselves. To this end we subjected leaves to mechanical wounding by forceps as well as applied a well-known plant immunity elicitor - bacterial flagellin (*flg22*). Since *flg22* and wounding have been well documented to induce different defence responses in leaves of a model plant *Arabidopsis thaliana*^11^ and are required in combination to activate defence in plant roots^12^, we also decided to test the combination of both wounding and *flg22*. This would mimic a natural scenario where bacterial infections follow mechanical plant damage by leaf herbivores, for example. Additionally, we also infiltrated a mixture of live plant pathogenic bacteria with a broad host range (*Pantotea aglomerans, Pseudomonas syringae pv. tomato, Erwinia rhapontici, Dickeya chrysanthemi*) to test for their ability to trigger local responses in the infected leaves. We selected a range of known plant immunity genes as potential markers for local leaf responses, designed primers for their homologs in *Medicago* truncatula and performed rt-qPCR at 2h, 4h and 6h following the application of the abovementioned stimuli. Leaves infiltrated with *flg22* and leaves treated with the combination wounding and *flg22* (*WF*) displayed strong induction of *MtPAD4* already at 2h, while *MtVSP1* expression was strongly induced at 4h and 6h following *WF* treatment (**Supplemental Figure 1**).

24h after stimulation *M. truncatula*, we tested the performance of fungal pathogen *Botrytis cinerea* on systemic leaves distal from the stimulated leaves. The *WF* treatment reduced the area of necrotic zones under *Botrytis* inoculation site compared to water controls, characteristic of systemic acquired resistance (SAR) (**Supplemental Figure 2**). Together with marker gene expression data, the *WF* treatment appeared to produce early local leaf responses as well as systemic responses that contribute to altered pathogen performance in our experimental system. Therefore, this treatment was chosen as a sender plant trigger for evaluating receiver plant responses in the following experiments.

### CMN mediates receiver plant responses to stimulation of signal senders

We grew two *Medicago truncatula* seedlings in a single pot, each in a separate 30 µm nylon mesh pocket that allowed AMF hyphae and root exudates to pass through while preventing direct root contact. One of the plants in each pot was inoculated with *Rhizophagus irregularis* spore suspension for seven weeks to establish a CMN between the two plants. In the CMN disconnection treatment (henceforth “cut”), we severed the CMN between plants using a blunt knife. In the CMN “connected” treatment, we preserved CMN connectivity, leaving the space between plants uncut. However, to separate the effects of CMN disruption from mere hyphal damage, we disrupted the extraradical hyphal network at the far edge of the “connected” pot. This approach controlled for potential plant responses to wounded AMF hyphae, allowing us to isolate the effect of CMN continuity - a key aspect missing from previous studies^9,10^. Moreover, the nylon mesh pockets also protected plant roots from accidental wounding during the CMN perturbations. Next, we mechanically wounded and applied *flg22* to the sender leaves or treated with water as the control (**Figure 1A).**

**Figure 1.**
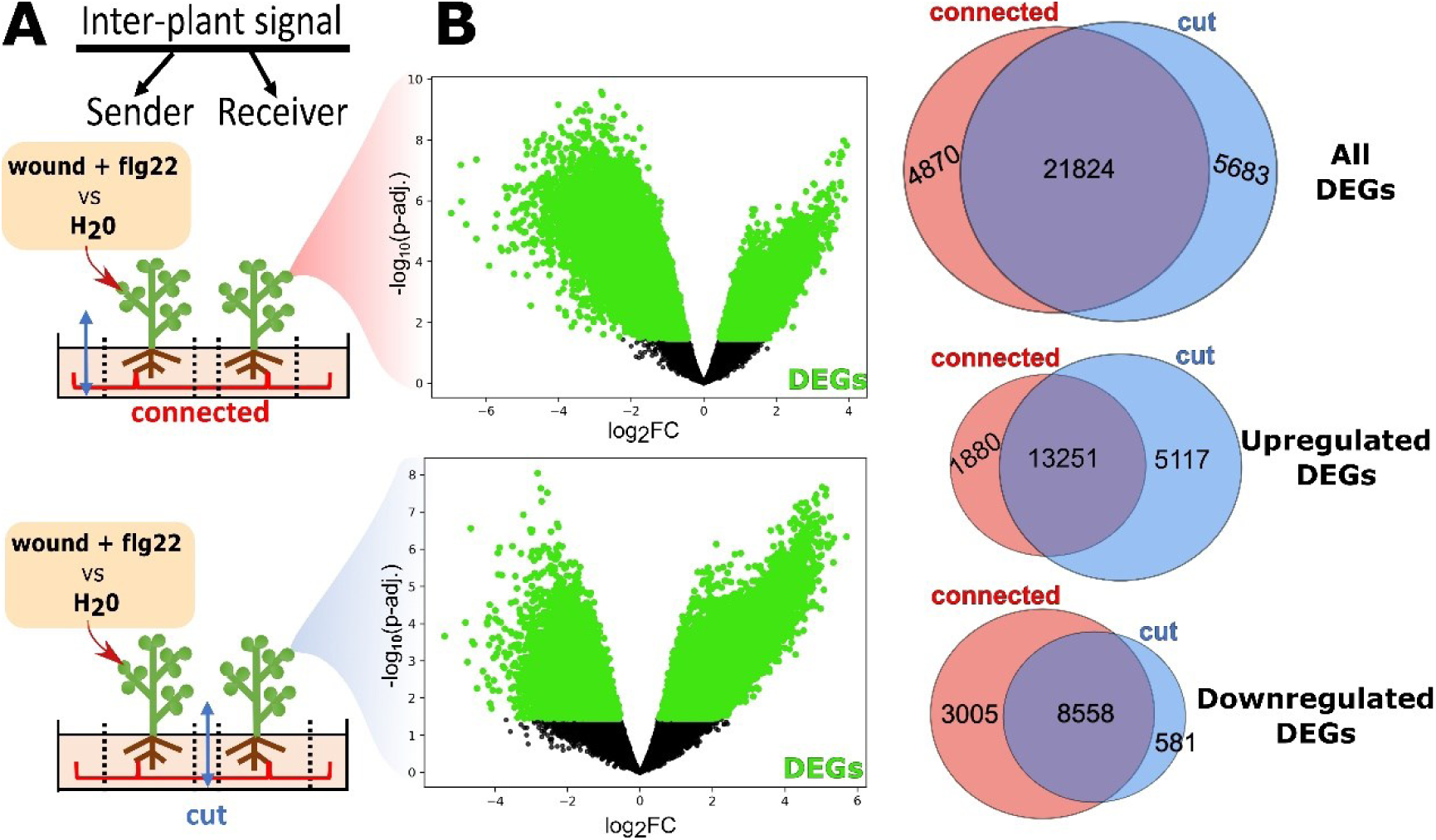
CMN connection contributes to wound and flagellin induced inter-plant responses. **(A)** A duet of *Medicago* plants were grown in 30 µm mesh pockets (dashed lines) and inoculated with AMF *Rhizophagus iregularis* to form CMN (red). Positions of mycorrhizal hyphal network cut are indicated with a blue double-head arrow. Sender plants with the cut-CMN and the CMN connection were stimulated by wounding and *flg22* application to measure responses in the receiver plant leaves. **(B)** Wounded and *flg22 (WF)* treated plants were firstly compared to corresponding H_2_O treatments in each CMN treatment respectively to identify up-and down-regulated responder plant DEGs. After, the DEGs in CMN connected pots (red circles) and the cut-CMN (blue circles) were compared. Volcano plot displays log10 of p-adj. and log_2_-fold change for each transcript in the *(WF)* comparison with water controls. Each treatment contained four biological replicates (N=4). Full list of DEGs, their annotations, fold change and FDR values are available in **Supplemental Table 1**.

To test whether there is any effect of the position of hyphal network perturbation without induced sender signals, we compared receiver responses to water treated senders in the pots with “cut” and “connected” CMN. Importantly, there was no significant transcriptomic change detected when the “cut” and “connected” treatments were compared (**Supplementary Figure 3**), strongly indicating that, firstly, the hyphal damage was likely applied evenly in both CMN treatments and, secondly, the position of the CMN interference alone does not contribute to any significantly differentially expressed genes (DEGs) in receiver plants. Therefore, this result permits a fair comparison how the integrity of inter-plant CMN connection contributes to unique receiver responses after sender stimulation by mechanical wounding and *flg22* application. To further test this, we stimulated the sender plants and compared receiver responses to non-stimulated senders in the “cut” and “connected” CMN treatments (**Figure 1)**.

We found that naïve receiver plants display distinct transcriptional responses to sender stimulation when CMN is cut compared to uncut inter-plant connection (**Figure 1B**). We identified 1880 up- and 3005 down-regulated DEGs that characterise CMN connection dependent receiver responses (**Figure 1B**). Among them, there were 15 DEGs that even showed opposite regulation in the cut and connected treatments (**Supplemental Table 1; tab-D)**. Twelve DEGs, including *M. truncatula* homolog of plant defence gene *PHATHOGENESIS RELATED 1* (MtrunR108HiC_038642), were significantly downregulated in the receiver plant connected by CMN but upregulated in disconnected CMN. In contrast, transcription factor NAC25 (MtrunR108HiC_019250) and a few receptor kinases such as peptidoglycan-binding LysM domain protein (MtrunR108HiC_035065) and RLK-Pelle-LRR-XII-1 family kinase (MtrunR108HiC_036113) were upregulated in the receivers with connected CMN but downregulated in the cut treatment.

### CMN-dependent inter-plant signals transcriptionally remodulate plant defence

Interestingly, gene functions relating to DNA metabolic processes - DNA repair (integration) and phosphotransferase activity - were significantly enriched within the 1880 upregulated DEGs (**Supplemental Figure 4; Supplemental Table 2 tab-A).** Functions with the most significant enrichment by downregulated DEGs were related to translation, RNA binding and processing, ribosomal subunit structure and organization as well as plant stress responses - primarily ROS metabolism, response to stimuli and jasmonic acid (**Supplemental Figure 5; Supplemental Table 2 tab-B).**

GO-term annotation was available for around 84% of the 4885 DEGs specific to the uninterrupted CMN treatment (**Supplemental Table 2 tab-C),** representing a good functional assessment. Complementary, we also performed an independent *de novo* functional prediction with all *M. truncatula* mRNA fasta sequences using *Mercator* ortholog-based function inference. Congruent with GO-term analysis, the *Mercator* enrichment analysis listed protein phosphorylation and phosphotransferase activity to be enriched by the upregulated DEGs **Supplemental Table 2 tab-D** and RNA processing and protein biosynthesis **Supplemental Table 2 tab-E** by the downregulated DEGs.

Intriguingly, after careful inspection of the original bin descriptors from the automated *Mercator* annotation (**Supplemental Table 3 tab-A)**, it was evident that many of the pre-assigned bins also relate to plant defence (**Supplemental Table 3 tab-B).** Therefore, we adopted a similar approach to the recent study in *Arabidopsis*^13^ and manually assigned a complementary “Defence signalling & response” functional bin (**Supplemental Table 3 tab-C)** for more accurate functional classification within the original hierarchy of *de novo* prediction by *Mercator* (**Supplemental Table 3 tab-D).**

Interestingly, the curated “Defence signalling & response” category contained higher cumulative transcript fold change of the 4885 DEGs for the CMN connected compared to the 5683 DEGs specific to the cut CMN treatment (**Figure 2A**) and displayed the strongest *WF* treatment induced inter-plant responses alongside protein biosynthesis and photosynthesis in receivers with inter-plant CMN connection (**Supplemental Table 3 tab-E)**. The defence signalling & response in CMN “connected” receivers showed downregulation of enzymes involved in ROS homeostasis, receptor-like kinases and intracellular MAPK and Ca-dependent kinases, as well as Ub-proteasome pathway (**Figure 2B**). Furthermore, genes involved in jasmonic acid synthesis and perception were downregulated compared to salicylic acid pathway. In addition, receiver plants displayed clear transcriptional changes in genes encoding specialized metabolites such as terpenoid and flavonoid synthesis.

**Figure 2.**
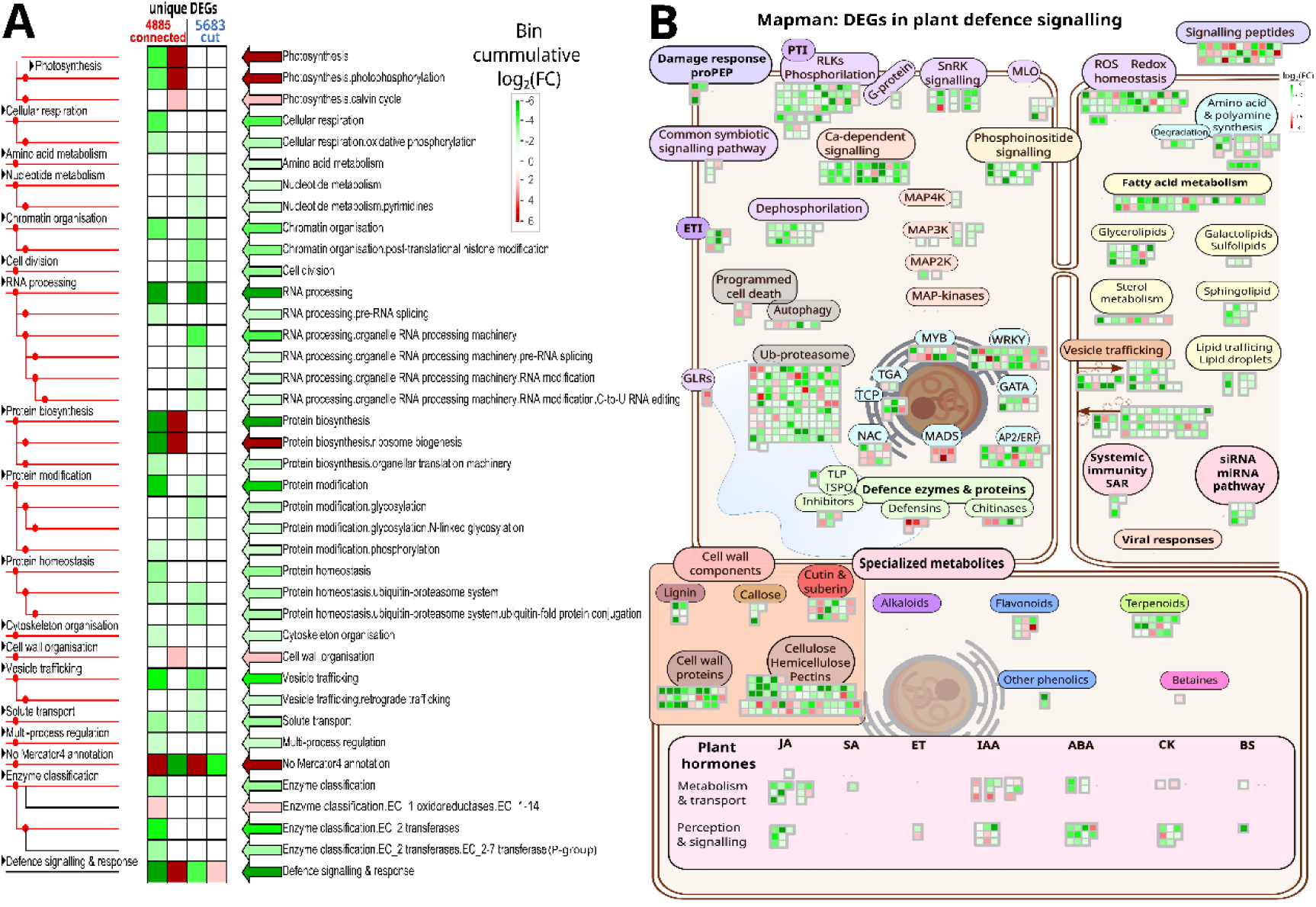
CMN connection is required for remodelling of biotic stress responses in inter-plant signal receivers. **(A)** Cumulative transcript fold changes of plant functions encoded by up- (red) and down-regulated (green) DEGs display different and specific responses to sender stimuli in the connected and cut CMN treatments. Visualisation is based on *Mercator4* functional annotation via *PageMan.* The ranked enrichment for each category is provided in **Supplemental Table 3 tab-E.** (**B**) 884 DEGs corresponding to plant biotic stress responses and plant defence signalling display dramatic transcriptional responses in responders with inter-plant CMN connection. Green colour represents downregulated, red – upregulated DEGs with colour intensity proportional to their log_2_FC. List of displayed DEG annotations and individual fold-changes is provided in **Supplemental Table 1 tab-B.** The image file for *MapMan* visualisation of “Defence signalling & response” pathways is available for download as **Supplemental figure 6**. Mapping file containing bin-assignment to immune receptors, intra-cellular signalling components and other defence related functions is available in **Supplemental Table 3 tab-D.**

An intriguing observation is the relatively high number of upregulated and low number of downregulated DEGs which are represent exclusive responses in the receivers with cut CMN compared to connected CMN (**Figure 1B**). Certain gene functions such as cell division, protein glycosylation, amino acid and nucleotide metabolism appear to be more enriched in the cut treatment (**Figure 2A**). Interestingly, when we investigated plant stress and defence related functions (**Supplemental Figure 7**), most of the transcripts from the cut CMN treatment displayed upregulation in sharp contrast to downregulation in the connected CMN treatment (**Figure 2B**). This strongly indicates that the receiver responses are highly influenced by the CMN integrity. Given that plant volatiles were not restricted neither in connected or cut treatment, this finding may suggest that the receiver responses to CMN-mediated signals could interact with responses from inter-plant volatile signals to affect gene regulation via yet unknown mechanisms.

### Inter-plant signals modulate volatile and metabolite composition in CMN-dependent manner

To further test whether inter-plant CMN connection influences the potential infochemical production and composition in receiver plant, leaf metabolites and volatiles were analysed using untargeted metabolomics approach. When *Medicago* senders were wounded and treated with *flg22* (**Figure 3A**), CMN-connected responders display significantly reduced production of volatile leucinamide (V2) and increased production of monoterpenoid β-fenchyl alcohol (V5) alongside several other leaf compounds such as medicagenic acids (L45; L47), acetylsoyasaponin (L46), genistein 7-O-glucoside (L15), 1H-Indole-3-acetaldehyde (L3), tyrosine (L8) and hex-bayogenin (L42) when compared to the H_2_O treated senders (**Figure 3B**). Notably, these compounds were not significantly over- or under-produced in inter-plant signal receivers with the interrupted CMN, suggesting CMN-dependent metabolic changes to neighbour signals. Instead, leaf metabolites soyasapogenol E (L51) and ethyl-D-glucopyranoside (L9) were significantly upregulated while mannopyranoside compound (L52), chrysoeriol (L19) and tricin GlcAGlcA (L20) were downregulated in receivers with the cut CMN (**Figure 3B**). Interestingly, receivers with the interrupted CMN did not show significant change in any measured volatile production unlike receivers with the CMN connection.

**Figure 3.**
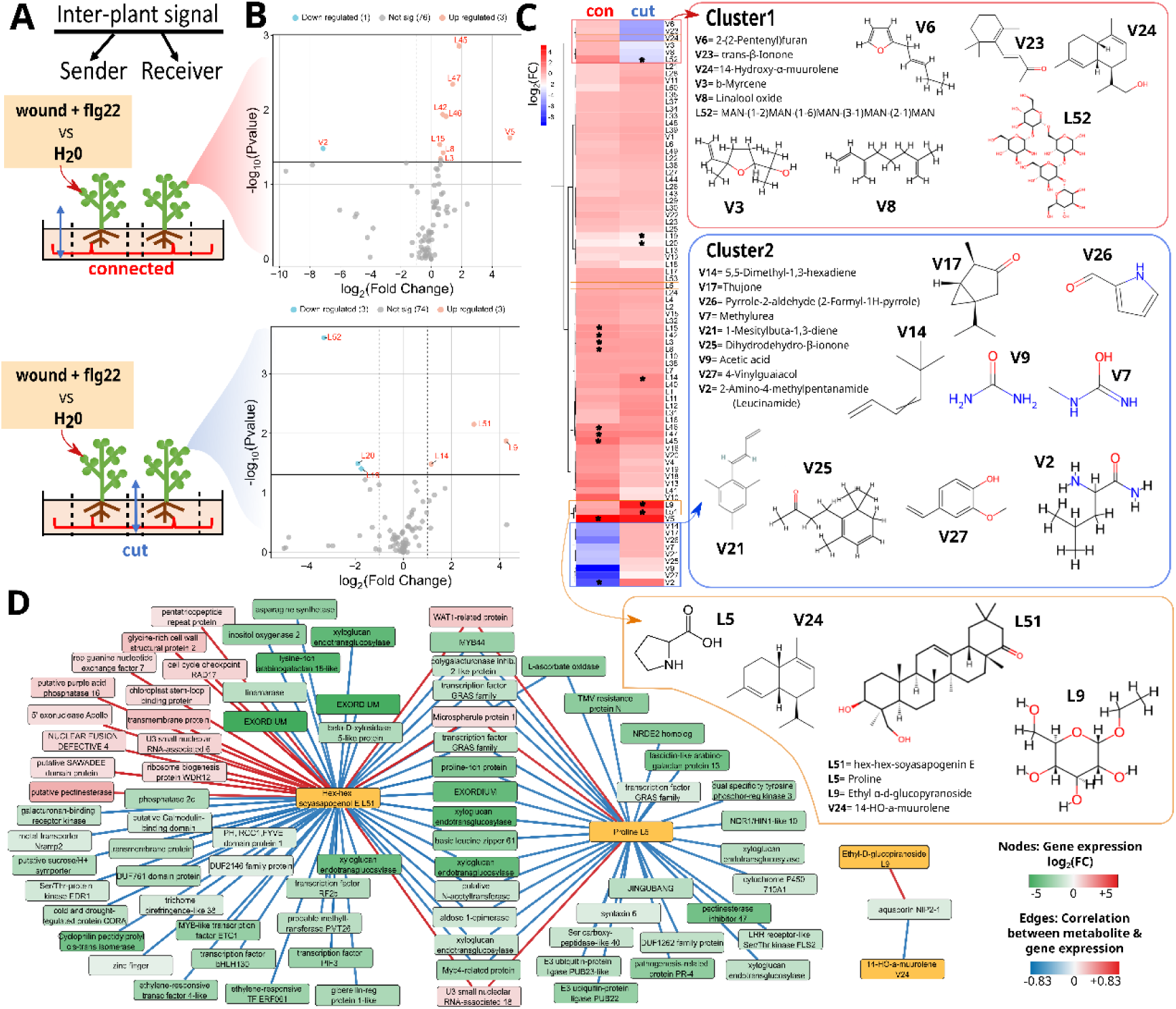
CMN modulates leaf metabolite and volatile production in response to neighbour stress-induced signals. **(A)** Two *Medicago* plants were grown in 30 µm mesh pockets (dashed lines) and inoculated with AMF *Rhizophagus iregularis* to form CMN (red). The blue double-head arrow indicates the position of CMN interruption prior to sender plant wounding and *flg22* application to measure responses in the receiver plant leaves. **(B)** Volcano plots displaying down-regulated or up-regulated leaf metabolites (L) and volatile compounds (V) when comparing inter-plant responses to wound and *flg22* with H_2_O controls in receivers with connected (top) or cut (bottom) CMN. **(C)** Hierarchical clustering of colour-coded log_2_ fold changes (FC) for all 53 detected leaf metabolites and 27 volatiles. Two clusters of compounds display the largest relative differences in receiver responses to inter-plant stress signals between connected (con) and cut CMN treatments. The asterisks indicate significant FC based on the volcano plots in panel **B**. Chemical structures are based on ChemSpider and PubChem databases for the closest matching compounds. **(D)** The network produced by regularized canonical correspondence (rCCA) analysis displays the correlation between normalized CPMS read counts of 4885 DEGs (identified in Fig. 1) with relative metabolite quantities. Only correlations with threshold >0.7 are displayed and include 4 metabolites (orange). The available homology-based gene function predictions from MtrunR108_HiC reference genome are mapped on the network along with colourcoded gene expression changes in CMN connected receivers. The source data for metabolite identification and network generation are provided in the **Supplemental Table 4.** Analytical standards were used to quantify selected metabolites and validate correspondence to independent samples in **Supplemental Figure 11**.

All leaf samples for metabolomics were collected from the same plants as the samples for RNA-seq, enabling investigation whether the is correlated change in leaf metabolites and defence gene expression using the set of 4885 DEGs specific to wounding and *flg22* response in CMN connected receivers. Surprisingly, proline (L5) and soyasapogenol E (L51) demonstrated the strongest correlation with a number of genes involved in plant defence responses (**Figure 3D),** including plant surface LRR Ser/Thr kinase receptors, including the flagellin receptor *FLS2*, *PATHOGENESIS RELATED 4,* E3 Ub ligases PUB22 and PUB23, ethylene and gibberellin response factors as well as several transcription factor families, especially MYB-related and GRAS. Additionally, aquaporin gene *NIP2* expression was strongly correlated with production of ethyl-D-gluccopiranoside (L9) and HO-α-muurolene (V24) (**Figure 3D)**.

### CMN mediated inter-plant signals induce acquired susceptibility to Botrytis cinerea but resistance to Fusarium sporotrichoides

Since we identified CMN-dependent changes in receiver plant biotic stress genes and leaf metabolites with reported functions in plant-pathogen interactions, we wished to test whether stressed sender plants modulate resistance to known plant pathogens in CMN connected receivers. We evaluated leaf infections by two plant generalist fungal pathogens - *Botrytis cinerea* and *Fusarium sporotrichoides*. Importantly, when the inter-plant CMN connection was cut, there was no significant difference in receiver lesion size upon *B. cinerea* or *F. sporotrichoides* infection between sender treatment of WF and water (**Figure 4B**). Interestingly, when we added generalist pathogen *F. sporotrichoides* to naïve receiver plants, the necrotic lesions were significantly smaller when sender plants were wounded and treated with flagellin (neighbour-primed) compared to water treated senders (**Figure 4A**), suggesting inter-plant signal induced resistance to the pathogen attack. However, when we exposed CMN connected naïve receiver plants to another pathogen *B. cinerea*, the necrotic lesions were significantly larger when sender plants were wounded and treated with flagellin compared to water treated senders (**Figure 4A**). Surprisingly, this effect was the opposite to intra-plant signals induced by wounding and flagellin treatment of distal leaves (SAR) which originally reduced lesion size in *B. cinerea* infected distal leaves (**Supplemental Figure 2**). This suggests that intra- and inter-plant signalling induced by wounding and flagellin may lead to different outcomes, possibly due to distinct underlying mechanisms. Our findings underscore the important role of the intra-plant CMN connection in how stress-induced signals affect pathogen resistance in neighbouring plants. Additionally, we highlight that stress-induced inter-plant signals in mycorrhiza-connected plants can result in varying levels of tolerance to different fungal pathogens in the receivers.

**Figure 4.**
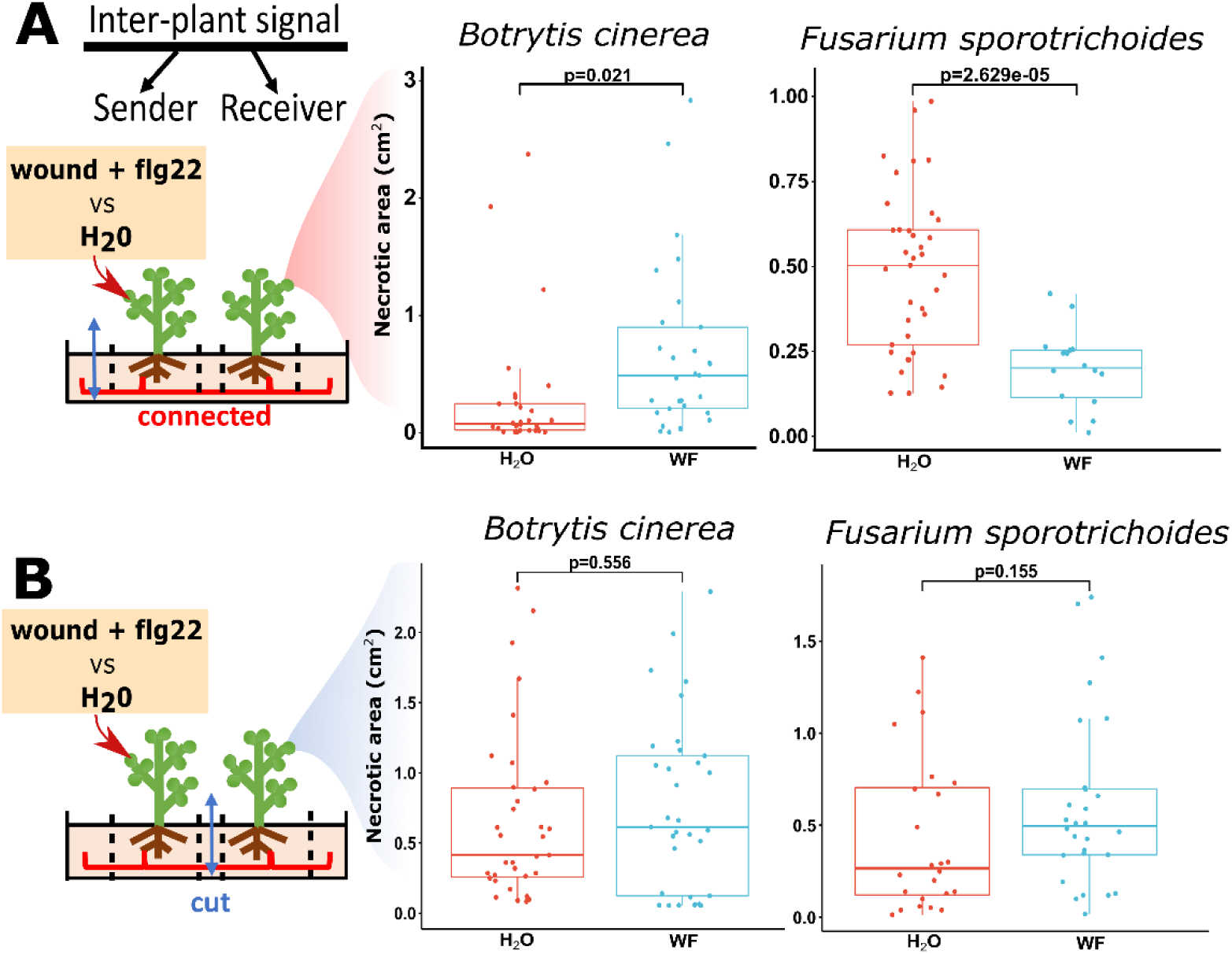
Stress-induced inter-plant signals modulate resistance to fungal pathogens in CMN connected plants. **A.** Size of necrotic lesions on *Mediago truncatula* leaves 72h after inoculation with *Botrytis cinerea* and *Fusarium sporotichoides* in CMN-connected plants. Wounding and flagellin (WF) treatment of sender plants induces resistance to *F. sporotrichoides* (F=21.355; p=2.629e-05) but susceptibility to *B. cinerea* (F=5.6603; p=0.02066). **B.** Size of necrotic lesions on *Mediago truncatula* leaves 72h after inoculation with *Botrytis cinerea* and *Fusarium sporotichoides* in receivers with cut CMN. WF treatment of sender plants does not significantly alter receiver plant lesion size upon *F. sporotrichoides* (F=2.089; p=0.1545) or *B. cinerea* (F=0.3496; p=0.5563) infection compared to receivers next to the H_2_O treated senders. Experiments in panel A and B were performed independent from each other. Pathogen assays in each graph were repeated independently 3-times and represented as pooled data. Line indicates the median, boxes represent the interquartile range (IQR), whiskers - variance within 1.5xIQR. Detailed comparison of all 3 individual experiments for each panel along with normality tests and repeated measures analysis is provided in **Supplemental figure 8**. Each treatment contained ≥8 biological replicates (receiver plants) per experiment.

## Discussion

### CMN as a mediator for inter-plant information exchange

For a few decades, CMN has been hypothesised to function as an inter-plant signalling conduit^3,4,14^. However, to date our mechanistic understanding on the molecular responses to stress induced signals in mycorrhizae-interconnected plants has been limited by very few empirical studies. Importantly, the interruption of CMN in tomato^9^ or pea^10^ plants was confounded by physical damage to symbiotic hyphae. Moreover, the few existing studies were all based on the targeted gene expression or enzyme activity measurements^9,15^, rather than characterising global responses of receiver plants to reveal the key pathways involved in the CMN-mediated neighbour priming against pathogens in an unbiased way. In this study, our experimental design allowed us to separate the effects of CMN disconnection from damage to the symbiotic hyphal network (**Supplemental Figure 3**). While *in vitro* studies have reported the capacity for reestablishment of hyphal connections after hyphal cutting^16^, the rate of hyphal growth, estimated to be around 10-20µm/h in vitro^17^ or 1-3mm/d on soil^18^ would not exceed the distance between nylon mesh in our experiment and unlikely to confound the experimental results.

For the first time, using an untargeted approach, we have identified a unique profile of volatiles, leaf metabolites, and transcriptional responses in receiver plants that share an AMF connection with biotic stress-stimulated signal senders, compared to plants with severed AMF connections (**Fig. 1**, **Fig. 3**). Furthermore, this manifested in altered plant susceptibility to pathogenic fungi *Botrytis cinerea* and *Fusarium sporotrichoides* **(Fig.4**). Together our study implicates CMN as important players for defence signal exchange. To establish this as a general phenomenon within plant communities in nature and agricultural systems, future research should expand the experimental scope to include diverse AMF and plant taxa and investigate the communality of plant regulatory and metabolic pathways that are modulated by neighbour signals. Furthermore, extended effects of CMN-mediated inter-plant signals to nodulation and plant interaction with plant growth promoting bacteria is another important avenue for future research in legume and non-legume crops and ultimate harnessing of CMN functions for agriculture.

### CMN contribution to modulation of inter-plant defence and metabolic responses

The majority of receiver leaf compounds responding to CMN-mediated neighbour signals are implicated in plant defence during biotic stress. One of the important key metabolic changes highlighted was the increased production of triterpenoid saponins (including hex-bayogenin (L42), medicagenic acids and their derivatives L45-L47) (**Fig. 3B**). Interestingly, medicagenic acids, bayogenin and other saponins are well-known important legume triterpenoids that act as phytoanticipins (preformed defence compounds) and phytoalexins (induced defence compounds), contributing to defences against pathogens by perturbing microbial membrane integrity ^19–23^. Additionally, levels of numerous monoterpene leaf volatiles were among the most notable differences between receivers in the connected and cut treatments (**Fig. 3C**). Monoterpenoid α-muurolene along with other terpenoids from endophytes of *Catharanthus roseus* are known to display antifungal properties^24^. These compounds display interplant stress-induced production in CMN connected receivers but not in the cut receivers.

Interestingly, receiver plants with intact AMF connection displayed increased production of 1H-Indole-3-acetaldehyde (IAAld) (L3) – an intermediate for auxin synthesis. As reviewed^25^ auxins have important roles in plant-microbe interactions. Curiously, *P*. *syringae* strain DC3000 produces IAA as a plant virulence factor from IAAld using dehydrogenase AldA in order to supress salicylic acid-mediated defenses in *A*. *thaliana*^26^. Furthermore, many auxin efflux proteins were significantly upregulated in the connected CMN treatment (**Supplemental Table 5 tab-A)**. Among them was vacuolar auxin transporter *WALLS ARE THIN1 (WAT1)-*related gene *MtrunR108HiC_014019. AtWAT1* is also known to be involved in cell wall thickening ^27^. Since CMN connected plants also displayed transcriptional changes in cell wall components as well as auxin synthesis and perception **(Fig. 2)**, we hypothesise that auxin homeostasis may in part contribute to cell wall remodelling and specialized metabolite production in defence against pathogens in CMN connected *Medicago* plants.

The integrity of CMN was not only responsible for increased metabolite production but also suppressed specific metabolites in response to neighbour signals. Certain triterpenes such as ethyl-D-gluccopiranoside (L9) or soyasapogenol E (L51) demonstrated more pronounced induction in the cut CMN treatment as opposed to the connected treatment, suggesting that CMN mediated inter-plant signals may dampen the inter-plant effects of other CMN independent signals. Soyasapogenol E levels were negatively correlated with expression of several ethylene response factors and members of MYB and GRAS transcription factor families (**Figure 3D)** which were well represented among the 4885 DEGs specific to the connected treatment (**Supplemental Table 6 tab-A)** and may indicate their regulatory roles in triterpenoid production. CYP450s and UDP-glycosyltransferases (UGTs) are primarily involved in mevalonate-dependent isoprenoid biosynthesis, including monoterpenes, sesquiterpenes and triterpenes^28^. We found sets of up- and down-regulated CYP450 and UGT enzyme transcripts that are specific to the cut and connected treatments (**Supplemental Table 6 tab-B)** and could be involved in the observed CMN-dependent differences in metabolite production. Lastly, leaf proline content was significantly correlated with expression patterns of several genes involved in plant defences, including *flg22* receptor *FLS2* and several E3 Ub ligases (**Figure 3D)**. Proline and its derivatives pyrroline-5-carboxylate are involved in plant responses to abiotic and biotic stress, including pathogens defence and mitigation of oxidative damage by scavenging reactive oxygen species (ROS)^29,30^. While the relative changes in proline content after sender stimulation were not significantly different between the cut and connected treatments (**Figure 3C)**, genes involved in ROS production and scavenging were significantly enriched among the 4885 DEGs specific to the connected treatment **(Supplemental Figure 5)**.

### Ecological and evolutionary roles of CMN-mediated inter-plant signals

CMN transmitted inter-plant stress signals have been previously considered to provide fitness benefits the receiver by providing increased pathogen resistance ^9^ and recruitment of natural enemies^10^. However, our data on differential inter-plant *M. truncatula* responses to *B. cinerea* and *F. sporotrichoides* infection clearly demonstrated that the outcome may not always be positive for the receiver. We emphasise that the potential beneficial effects from eavesdropping on conspecific signals via CMN must be holistically studied in context of different pest and pathogen species as well as host genotypes. Intriguingly, neighbour-modulated susceptibility (NMS) has been documented in rice and durum wheat growing next to pathogen free conspecifics of different genotypes and appears to be mediated by inter-plant signals in soil ^31^. In particular, the study showed that wheat growing next to a different conspecific genotype display increased susceptibility to hemibiotroph *Zymenoseptoria tritici* but reduced susceptibility to biotroph *Puccinia triticina,* suggesting different production or perception of signals by different plant genotypes. Similarly, *Magnoporthe oryzae* showed fewer lesions on rice neighbouring a different conspecific genotype compared to isogenic cultivar, while another hemibiotroph *Bipolaris oryzae* demonstrated more lesions ^31^. *Arabidopsis* plants exposed to elicitors in insect eggs are able to produce inter-plant signals that induce broad spectrum resistance against bacterial pathogen *Pseudomonas syringae* ^1^ and fungus *Botrytis cinerea* ^32^, suggesting that certain plant-to-plant signals may be potentially more conserved and less specific to different pathogen species. This further highlights the need to test a range of different elicitors on senders and evaluate responses in responders of different genotypes and species in the future experiments.

However, in our foundational experiments presented herein we wished to use stimuli or elicitors that would give consistent and reproducible PAMP or DAMP responses without the added complexity and potential variability of using live herbivore or pathogen infections. For example, insect oral secretions are known to contain additional chemical elicitors that are recognised by plant HAMP receptors or deliver insect effector proteins that supress plant defence responses^33^. Similarity, microbial pathogens produce effectors that either supress PTI responses in plants or induce effector-triggered immunity (ETI)^34^, thus introducing mechanistic complexity for the generation of systemic and inter-plant signals in the senders. Furthermore, different feeding guilds of herbivores, such as cell-content feeding, leaf or root chewing, stem or leaf boring, phloem or xylem sucking, as well as the range of pathogens with biotrophic to necrotrophic lifestyles and diverse effector repertoires offer an extended scope for future investigations on plant-to-plant responses to these diverse interactions.

Neighbour induced susceptibility or resistance would provide opposing selective pressures for the evolution of specific signal generation and perception mechanisms along with receiver responses. For example, volatiles are known for their roles in pollinator as well as parasitoid attraction. Since *Medicago truncatula* is primarily a self-pollinator, it is unlikely that the altered receiver VOC profiles would affect plant reproductive fitness directly but may be important in other outcrossing species. Interestingly, volatile monoterpenes (e.g., penene, campene, sabinene) have been shown to induce inter-plant systemic acquired resistance (SAR) against *Pseudomonas*^2^ and could potentially relay SAR signals from plant-to-plant^35^. While we did not find changes in the same monoterpene compounds^2^ it would be intriguing to test whether VOCs could further propagate and amplify CMN-mediated inter-plant signals and thus affect plant defences at a community level. Moreover, VOC and specialized metabolite composition is very diverse across angiosperms. For example, the lineage-specific number and expansions of terpene synthase genes across angiosperms suggest functional divergence in terpene structure and regulation in different ecological conditions and plant genotyopes^36–38^. Therefore, metabolic effects of CMN-mediated inter-plant stress signals could have diverse effects in crops under field conditions.

## Conclusion

Receiver plants’ biotic stress responses and defence signalling pathways can be uniquely shaped by inter-plant signals transmitted via CMN and manifest in differential disease outcomes. These findings underscore the potential pivotal role of CMN in modulating plant immunity at a community scale, revealing a sophisticated and nuanced mechanism of pathogen-specific susceptibility and resistance. This insight prompts further investigation into the mechanisms of CMN-mediated interplant signalling, which could be transformative for understanding plant defence dynamics in complex natural ecosystems and for optimizing disease management strategies in agriculture.

## Materials and Methods

### Plant growth

30 µm pore size autoclavable PA6-grade nylon mesh (JPP140 from Technical textiles, Ekobalta, Latvia) was used to make 11 cm wide and 10 cm deep pockets which were filled with autoclaved soil:sand mixture (3:1 v/v). Soil (gardening substrate with black earth) was purchased from Biolan Baltic OÜ (pH: 6,0; EC: 25 mS/m; N∼100 mg/L; P∼60 mg/L; K∼200 mg/L; peat fraction<20mm). Quartz sand (grain size ≤0.5mm) obtained from SACRET^®^, Latvia. Two substrate filled mesh pockets were placed into 1L black plastic pot (11×11×12 cm L x W x H) and surrounded with the surplus soil:sand mixture to prevent them from touching each other or the edge of the pot. *Medicago truncatula* R108 seeds were scarified with a sterile scalpel and germinated on wet *Whatman* filter paper for 2-3 days before planting onto the soil:sand mixture within each mesh pocket. The images of the setup are provided in the **Supplemental Figure 9**. Plants were grown in growth cabinet (SANYO MLR-351H, Japan) at 20℃, 50%RH, 16h day / 8h night cycle using a set of twelve 6500K (3200lm) and three 2700K (3350lm) fluorescent lightbulbs (T8 MASTER TLD G13 Philips, 36W) to cover the entire PAR spectrum with 300 mmol m^2^ s^−1^ light intensity. Each pot was watered with 200mL 0.5X Hoagland nutrient solution once a week.

### AMF cultivation & inoculation

*Rhizophagus irregularis* (DAOM 197198) was cultivated on *Daucus carota* hairy root cultures ^39–41^, grown on M-media (1% sucrose, 0.4% Gelrite gellan gum) at 25℃ in darkness. 6-month-old cultures were processed for *R. irregularis* spore collection using 10mM citrate buffer (pH=6) to dissolve spore-containing solid media and capture spores within 40µm cell stainer (SPL Lifesciences) and resuspend in sterile dH_2_O. Spore density was determined microscopically. Suspension volume containing 200 spores was applied to one of the seedlings in the pot. Plants were grown for 7 weeks to establish CMN between the sender and receiver plants. The colonization status of the receiver plant was verified using ink-vinegar staining method ^42^ after the experiment.

### Treatment of CMN and sender plants

7-weeks post AMF inoculation all experimental pots were divided into two groups: 1) uninterrupted CMN (sender-receiver plants remain connected via CMN) where a blunt knife was inserted into the soil along the edge of the pot as a control and 2) a group of cut CMN where a blunt knife was inserted into the soil between the mesh pockets to disrupt the CMN connection between signal sender and receiver plants as depicted in **Supplemental Figure 9**. Three days after this manipulation, sender *Medicago* plants were subjected to mechanical wounding by applying three squeezes with metal forceps to each leaflet of two fully expanded trifoliate compound leaves, followed by the application of a 5 μL drop of synthetic 1 μM bacterial flagellin (*flg22*, MedChemExpress, Sweden) on each forceps-wounded leaflet. 5 μL of dH_2_O without mechanical wounding was used as the control treatment. Receiver plants were left untreated and collected 3 days after the stimulation of the sender plants.

### RNA extraction & transcriptome sequencing

Three separate trifoliate leaves from 8-weeks old receiver plants were harvested in liquid N_2_, homogenized with pestle and mortar for RNA extraction using ReliaPrep^TM^ RNA Miniprep System (Promega, cat. Z6012). RNA quantification was performed with Qubit™ RNA BR Assay Kit and Agilent Bioanalyzer RNA 6000 Pico Kit. Samples from 4 different plants were processed as biological replicates for each treatment. Total 400ng of each RNA sample (RIN≥7, OD_260/280_ between 1.9-2.1, OD_260/230_ between 1.5-2.0) was processed for library preparation using MGIEasy RNA Directional Library Prep Set following manufacturer’s instructions. QIAseq FastSelect –rRNA Plant Kit (QIAGEN) was used to deplete rRNA prior to the library fragmentation step. Qubit^®^ dsDNA HS Assay and Agilent HS DNA Kit was used to determine library concentration and length (average 326 nt) prior to multiplexing and circularization. The final pooled libraries (1.94 ng/µL) were sequenced on DNB G400RS high throughput sequencing machine (MGI) using FLC PE150 flowcell.

### Raw data processing and DEG analysis

Read filtering and adapter removal was performed with *fastp 0.20.0* (default parameters; read trimming >100 bp). *STAR 2.7.10a* ^43^ was used to map the reads against *Medicago truncatula R108* reference genome available in https://medicago.toulouse.inrae.fr/MtrunR108_HiC (INRA, Toulouse). *SortMeRNA version 4.3.6* ^44^ was used to quantify and filter rRNS using smr_v4.3_default_db database. DEG analysis was performed with *limma* (Bioconductor)^45^. Low count features were filtered using the function *filterByExpr* implemented in *edgeR* (v4.6.2). *Voom* with quality weights was used to transform the data prior to linear modelling with *limma*. Features with FDR < 0.05 were regarded as statistically significant and used for downstream selection of CMN connection specific interplant responses.

### Functional enrichment analysis

Enrichment of GO biological and molecular functions for DEGs was calculated using *agriGO v2.0* ^46^ Fisher exact test (FDR) with *MtrunR108_HiC_functional annotation* file as reference and visualized with enrichment bubble *SRplots*. To independently validate our enrichment analysis, we performed complementary enrichment analysis with the *de novo* gene function predictions obtained with *Mercator4 v2.0* tool^47^. To this end, *MtrunR108_HiC.1.0.fasta_features* (all genomic mRNA nt sequences) were used as input for the gene orthology search (**Suppl. Table 3; tab-A**). The automated function descriptors were manually checked and complementary assigned to “Defence signalling and response” bin (**Suppl. Table 3; tab-B**). Since a distinct and comprehensive “plant stress and defence” function category was originally absent from the automated functional assignment (**Suppl. Table 3; tab-A**), we extracted the corresponding gene identifiers, and *de novo* compiled the ‘plant stress and defence” bin (BINCODE 80) (**Suppl. Table 3; tab-C**). This list was then combined with the original list of Mercator4 functions from **tab-A** to produce an updated input (mapping) file for functional enrichment analysis, available in **Suppl. Table 3; tab-D**. Subsequent visualisation of DEG functions was performed with *MapMan v3.7* ^48^ and *PageMan* ^49^ functional annotation tools. Statistical analysis on DEG enrichment for *MapMan* pathways was performed using built-in Wilcoxon rank test and Benjamini-Hochberg (BH) p-value correction^50^. The graphical overview of plant defence pathways displayed in **Fig. 2** was drawn manually in *Inkscape* and loaded as separate pathway image file into *MapMan* (available for public download as .jpeg in **Figure supplement 6** and reusable by citing this manuscript).

### rt-qPCR

To verify the expression data from RNA-seq dataset, we performed rt-qPCR on selected genes from independent samples. Test genes were selected based on the largest FC difference among connected and cut treatments (**Supplemental Figure 10A**). Reference gene candidates were selected based on the lowest fold change across all treatments (**Supplemental Figure 10B**). *M. truncatula* gene specific primers were designed in NCBI Primer BLAST for 100-200 nt amplicons spanning exon-intron junction where possible. cDNA was synthesised with *Maxima First Strand cDNA Synthesis Kit (Thermo Fisher Scientific)* and qPCR performed with *Maxima SYBR Green/ROX qPCR Master Mix (2X) (Thermo Fisher Scientific)* on *ViiA™ 7 Real-Time PCR System (Applied Biosystems)* and analyzed with the native *QuantStudio 7 Pro 1.6.1* software. Primer efficiency (E%) was calculated using the formula 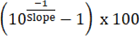, with slope being obtained by plotting the log_10_ values of 10-fold serial dilutions from 1ng to 1ag of purified cDNA (**Supplemental Figure 10E**). The expression of each test gene was normalized by the geometric mean of two selected reference genes (*MtEXO70* and *MtBP28*) using the formula relative expression =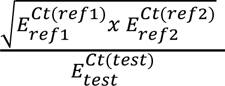, where *E_ref_*– primer efficiency of reference gene, *E_test_* – primer efficiency of test gene, *Ct(ref)* – Ct value of reference genes, *Ct(test)* – Ct value of the test gene.

### Leaf metabolic analysis with LC-HRMS

Metabolome was analysed from the same leaf samples as plant transcriptome by aliquoting 200mg of the frozen homogenized leaf tissues and resuspending in 1.5 mL of 70% methanol, vortexed for 1 minute and left for maceration for 5 days at 4°C in darkness. The obtained extract was filtered through a 0.45 µm filter and injected into the Agilent 1290 Infinity II series HPLC system combined with Agilent 6530 qTOF MS system (Agilent Technologies, Deutschland GmbH, Waldbronn, Germany). Zorbax Eclipse Plus C18 Rapid Resolution HD (2.1×150mm, 1.8μm particle size) column was used at a flow rate of 0.3 mL/min. The column oven was set at 45°C, and the sample injection volume was 3 μL with 40 sec needle wash (70% methanol). The mobile phase consisted of a combination of A (0.1% formic acid in water) and B (0.1% formic acid in acetonitrile). Gradient elution program was used as follows: Initial 5%B, 0-6 min 20%B, 6-20 min 40%B, 20-23 min 80%B, 23-28 min 95%B, 28-33 min 95%B, 33-39 min 5%B. UV/VIS spectra were recorded at 280 nm and 330 nm. The adjusted operating parameters of the mass spectrometer were set as follows: Fragmentation: 70 V; Gas temp: 325°C; Drying gas 10 L/min; Nebulizer 20 psi; Sheath gas temp 400°C; Sheath gas flow 12 L/min. Electrospray ionization (ESI) was used as a source in operating in positive mode. The internal reference masses of 121.050873 m/z and 922.009798 m/z (G1969-85001 ES-TOF Reference Mass Solution Kit, Agilent Technologies & Supelco) were used for all analyses of the samples. The Agilent MassHunter Qualitative analysis 10.0 data acquisition software was applied to analyze LCMS data and compare ion fragmentation data with information available in literature and databases, such as METLIN, LIPID MAPS®, Phenol – Explorer, Lotus - Natural 575 Products Online and FooDB v1.0 with acceptance mass error ≤ 20 ppm and consistency of isotope/adduct patterns.

Analytical standards (L-proline, jasmonoyl-isoleucine, medicagenic acid, apigenin, soyasapogenol B) were used for absolute quantification of matching compounds and their derivatives. External calibration was confirmed at MSI Level 1 by co-injection RT match (ΔRT ≤ 3%) and HR m/z for the dominant ESI(+) adduct (± 5 ppm). Quantification used external solvent-based calibration (7 levels; 3 replicates per level; linear fit with 1/x weighting when required). No internal standards or matrix-matched/standard-addition approaches were applied. Primary stock solutions were prepared at approx. concentration 0.1 mg mL⁻¹ in methanol except jasmonoyl-isoleucine in ethanol, then serially diluted in methanol to the working calibration levels. Validation data are summarised in **Supplementary Table 4 tab-B**. Measured concentrations (µg/g FW) with independent samples are reported in **Supplementary Table 4 tab-C.** The correspondence of the repeated experiment and the experiment reported in Figure 3 is displayed in **Supplemental Figure 11**.

### Plant volatile analysis

Volatile compound analysis was performed using the same leaf samples as the plant transcriptome analysis. The samples were ground into a fine powder using liquid nitrogen, and 100 mg of the homogenized leaf tissue was aliquoted into clean 20 mL headspace vials. A saturated NaCl solution was added to prevent enzyme reactions, and the vials were sealed with Agilent headspace silica gel caps. Each vial was then heated for 20 minutes at 60 °C with an agitator cycle of 30 seconds on and 15 seconds off. Samples were injected into an Agilent Technologies 7820A gas chromatograph coupled with an Agilent 5977B mass selective detector (MSD) using a Gerstel MPS Autosampler. The syringe temperature was set to 130 °C, and the injection volume was 2500 μL. A polar CP-Wax 52CB capillary column (50 m × 0.32 mm, 0.20 µm film thickness) with polyethylene glycol was used. Helium (He) served as the carrier gas with a split ratio of 1:20 and a flow rate of 1.2 mL/min. The temperature program started at 60 °C, increased at a rate of 10 °C/min to 250 °C, and was held for 3 minutes. The injector temperature was set at 260 °C. Mass spectra were recorded at 70 eV within an m/z range of 50–500. The ion source temperature was maintained at 230 °C. Compound identification was based on retention indices (determined using a homologous series of C5–C24 n-alkanes) and mass spectral comparison with the NIST (National Institute of Standards and Technology) MS Search 2.2 library. Data acquisition and analysis were performed using Agilent MassHunter Qualitative Analysis 10.0 software.

Target volatiles (β-myrcene, linalool, β-fenchyl alcohol) were confirmed at MSI Level 1 by retention index (vs C5-C24 n-alkanes) and EI spectral matches to authentic standards. Primary stocks (1 mg mL⁻¹ in methanol) were used to prepare calibration standards: each 20 mL HS vial contained 4.7 mL of 30% (w/v) NaCl and 0.3 mL of the appropriate standard solution; working levels were obtained by serial dilution to five concentrations (100–3000 ng mL⁻¹) with three replicate vials per level. Quantification used external calibration (linear regression; 1/x weighting when required). Validation outputs are reported in **Supplementary Table 4 tab-E**.

### Correlation of gene expression and metabolite data

Regularized canonical correspondence analysis (rCCA) was used with the *mixOmics* R package (available from Bioconductor; http://www.mixOmics.org) in R studio. R script provided in **Supplementary Table 4 tab-D.** Correlations >0.7 were visualised via *Cytoscape* v3.10.3.

### Plant pathogen assays

*Fusarium sporotrichoides (Fs)* and *Botrytis cinerea (Bc)* were grown on 0.5X PDA at 20°C in darkness for 4 weeks. Mycelial plugs, obtained with a 7.5mm diam. cork borer, were placed on three different trifoliate leaves. Each leaf was placed in a 5.5cm diam. Petri dish with water-saturated filter paper to maintain moisture level. The edge of the petri dish had a pre-cut slit for the leaf petiole to remain attached to the plant throughout the experiment. Plants were placed in shaded at 20°C for 72h. Agar plugs were removed before leaf photography. *ImageJ (Fiji)* was used to measure the area of necrotic lesions.

## Data availability

RNA-seq raw data (fastq) are uploaded to NCBI under BioProject: **PRJNA1186668**.

## Author contributions (Contributor Role Taxonomy CRediT)

**ZO:** conceptualization and experimental design (1), data curation (2), formal analysis (3), funding acquisition (4), investigation (5), methodology (6), project administration (7), reagent acquisition and lab resource planning (8), student supervision (10), data validation (11), statistical analysis and visualisation (12), writing (13), reviewing and editing the original manuscript (14)

**IN**: investigation – metabolic analysis (5), reviewing the original manuscript (14)

**EV:** formal analysis (3) of pathogen data, investigation – plant-pathogen assays (5), reviewing the original manuscript (14)

**DP**: investigation – experiment setup, RNA extractions, rt-qPCR (5), reviewing the original manuscript (14)

**AK:** investigation – experiment setup, RNA extractions, rt-qPCR (5), reviewing the original manuscript (14)

**DS**: investigation – qPCR validation (5)

**AA:** investigation – root staining for AMF colonization (5)

**KTB**: investigation – experiment setup, RNA extractions (5), reviewing the original manuscript (14)

**IS**: data curation (2), formal analysis (3) of RNA-seq data, reviewing the original manuscript (14)

**SJL:** conceptualization and experimental design (1), methodology (6), reviewing and editing the original manuscript (14)

## Acknowledgements

We are grateful to Prof. Yves Poirier group (DBMV, University of Lausanne) for the generous donation of *Medicago truncatula* R108 seeds. We also thank BMC Genome Centre and Bioinformatics Unit support staff for assistance during sequencing and raw data handling.

## List of Supplemental tables

**Supplemental Table 1:** Differentially expressed receiver plant genes (DEGs) in response to sender stimulation by wounding and flg22 in cut and connected CMN treatments.

**Tab A**: log2(fold changes) and gene descriptions of 21 809 responder plant DEGs shared in the cut and connected CMN treatment.

**Tab B**: log2(fold changes) and gene descriptions of 4885 responder plant DEGs in connected CMN treatment.

**Tab C**: log2(fold changes) and gene descriptions of 5698 responder plant DEGs in cut CMN treatment.

**Tab D**: log2(fold changes) and gene descriptions of 15 responder plant DEGs which show opposite regulation in cut and connected CMN treatment.

**Supplemental Table 2:** GO-term enrichment for the 1880 upregulated and 3005 downregulated receiver plant DEGs in the connected CMN treatment. Functional enrichment is based on Fisher exact test with Hochberg (FDR) correction using agriGO v2.0. Significantly enriched functions (p<0.05; FDR<0.05) for each ontology category - biological process, molecular function and cell compartment - are highlighted in bold and coloured.

**Tab A**: GO enrichment for the 1880 upregulated DEGs displayed in Figure 1B. Supplemental Figure 4 displays sign. enriched Go-terms corresponding to biological function

**Tab B**: GO enrichment for the 3005 downregulated DEGs displayed in Figure 1B. Supplemental Figure 5 displays sign. enriched Go-terms corresponding to biological function and molecular function. Biological function GO terms related to stress response are manually highlighted in yellow

**Tab C**: GO-term representation (%) among up- & down-regulated responder plant DEGs and the Medicago truncatula R108 reference genome

**Tab D**: List of significantly enriched Mercator4 functional bins containing upregulated receiver DEGs in connected CMN treatment

**Tab E**: List of significantly enriched Mercator4 functional bins containing downregulated receiver DEGs in connected CMN treatment

**Supplemental Table 3:** assignment of plant stress and defence bin for functional enrichment analysis of CMN-mediated receiver plant DEGs

**Tab A**: original Mercator4 functional annotation of 39029 Medicago truncatula R108 genes based on sequence orthology from all mRNA FASTA files as input.

**Tab B**: descriptors of Mercator4 bins that correspond to Plant stress and signalling functions. Only the original Mercator4 output bins from tabA and their corresponding genes are considered. References from the published literature justify the the assignment of bin decriptors to “Defence signalling and response” bin.

**Tab C**: Mercator4 BIN contents that are reassigned for the “Defence signalling and response” bin (BINCODE 80)

**Tab D**: list of all original Mercator4 functional annotations (tabA) plus the manually reassined “Defence signalling and response” bin (BINCODE 80) from TabC. TabD=TabA+TabC. TabD was used for the enrichment analisis in TabE.

**Tab E**: Enrichment of Mercator4 functional categories, taking into account the cumulative fold-change of 4885 up-and down-regulated responder DEGs in connected CMN treatment. Analysis performed Mann Whitney U Test (Wilcoxon Rank Sum Test) and Benjamini-Hochberg p-value correction for multiple comparisons to reveal whether the average response (cumulative log2FC) of a BIN is different from the response of all the other BINS. The results are dynamically calculated upon loading of an experiment file with fold changes for a chosen gene list into MapMan tool.

**Tab F**: List of all 39029 Medicago truncatula R108 genes and subset of Mercator4 annotated functions within the “Defence signalling and response” bin (BINCODE 80), representing 16.9% of total transcripts

**Supplemental Table 4:** Source data for compound identification using LC-HRMS and HS-GC-MS as well as rCCA and correlation network between transcriptome and metabolome datasets.

**Tab A**: LC-HRMS data with tentative metabolite id & peak areas used in Figure 3

**Tab B**: LC-HRMS data calibration with selected standards

**Tab C**: LC-HRMS data for compound quantification with analytical standards in original & independent samples displayed in Supplemental Figure 8

**Tab D**: HS-GC-MS data with tentative metabolite id & peak area used in Figure 3

**Tab E**: HS-GC-MS data calibration with selected standards

**Tab F**: HS-GC-MS data for compound quantification with analytical standards

**Tab G**: log2(FC) of compound levels in receivers of WF stimulated vs H2O treated sender signals used to generate volcano plots and heatmap in Figure 3B&C

**Tab H**: data matrix input for rCCA analysis in Figure 3D; columns with gene and metabolite id; rows - treatments with log2 normalized gene read counts (limma package; CPM counts per million +1 pseudocount) and log2 normalized relative metabolite abundances

**Tab I:** raw network data obtained with rCCA; from. Nodes describe metabolites; to. nodes describe genes; gene id and corresponding descriptions based on Medicago truncatula R108 reference genome available in https://medicago.toulouse.inrae.fr/MtrunR108_HiC (INRA, Toulouse); FC values based on RNA-seq data wf_connected vs h2o_connected; edge weight is correlation strength based on rCCA for all nodes with correlation >0.7

**Tab J**: rCCA script in R for correlation network analysis to generate Figure 3D

**Supplemental Table 5.** Comparison of transcription factors, auxin transporters, P45 and UGT enzyme fold change in receiver from “connected” and “cut” treatments.

**Tab A:** log2(FC) of upregulated and downregulated DEGs for auxin transporters, auxin response factors, GRAS, MYB transcription factors as well as ethylene response factors in CMN connected receivers

**Tab B:** log2(FC) of DEGs corresponding to sender signal regulated UDP-glycosyltransferases and cytochrome P450s in “connected” and “cut” treatments

**Tab C**: log2(FC) of non-DEG (p-adj.>0.05) cytochrome P450 and UGT transcripts in the “connected” and “cut” treatments

## Supplemental figures

**Supplemental Figure 1.**
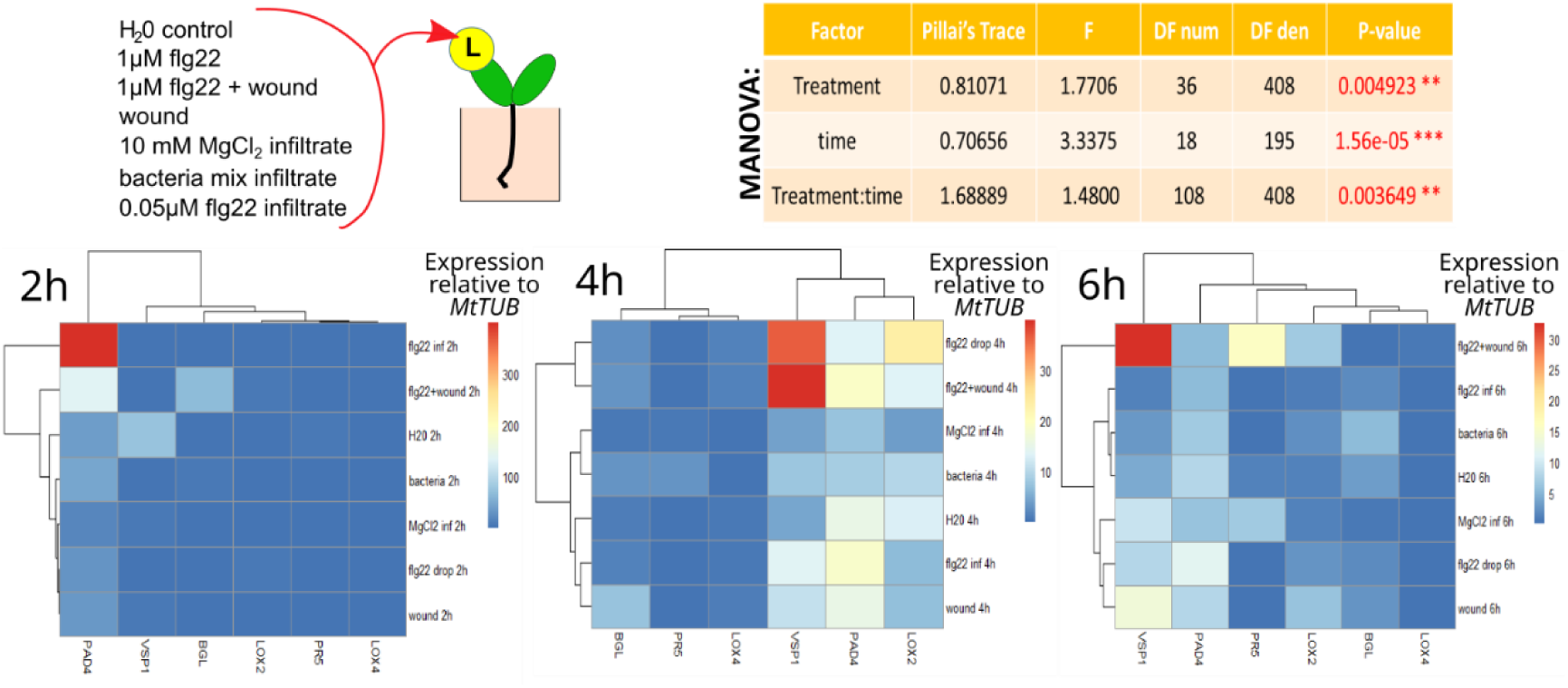
Local expression of *Medicago truncatula* immunity markers (*MtPAD4, MtVSP1, MtBGL, MtLOX2, MtLOX4, MtPR5*) relative to *MtTUB* reference gene 2h, 4h and 6h following the application of various stimuli on leaves: mechanical wounding by forceps (3x squeezing leaf blade), 5 µL drop of 1µM flg22, 5 µL drop of dH_2_O (control), syringe infiltration of 0.05µM flg22, bacterial mix in 10mM MgCl_2_ or sterile 10mM MgCl_2_ as the control. Bacterial mix consisted of live suspension of overnight LB cultures of *Pseudomonas syringae, Dickeya chrysanthemi, Erwinia rhapontici, Pantotea agglomerans* at OD_660_=0.0005 each. Heatmap displays average expression of 4 independent biological replicates. Each replicate was a pooled sample from two mature stimulated trifoliate leaves from separate plants. 2-factor Multiple Analysis of Variance (MANOVA) indicates significant changes in gene expression over the time-course and differences among the treatments.

**Supplemental Figure 2.**
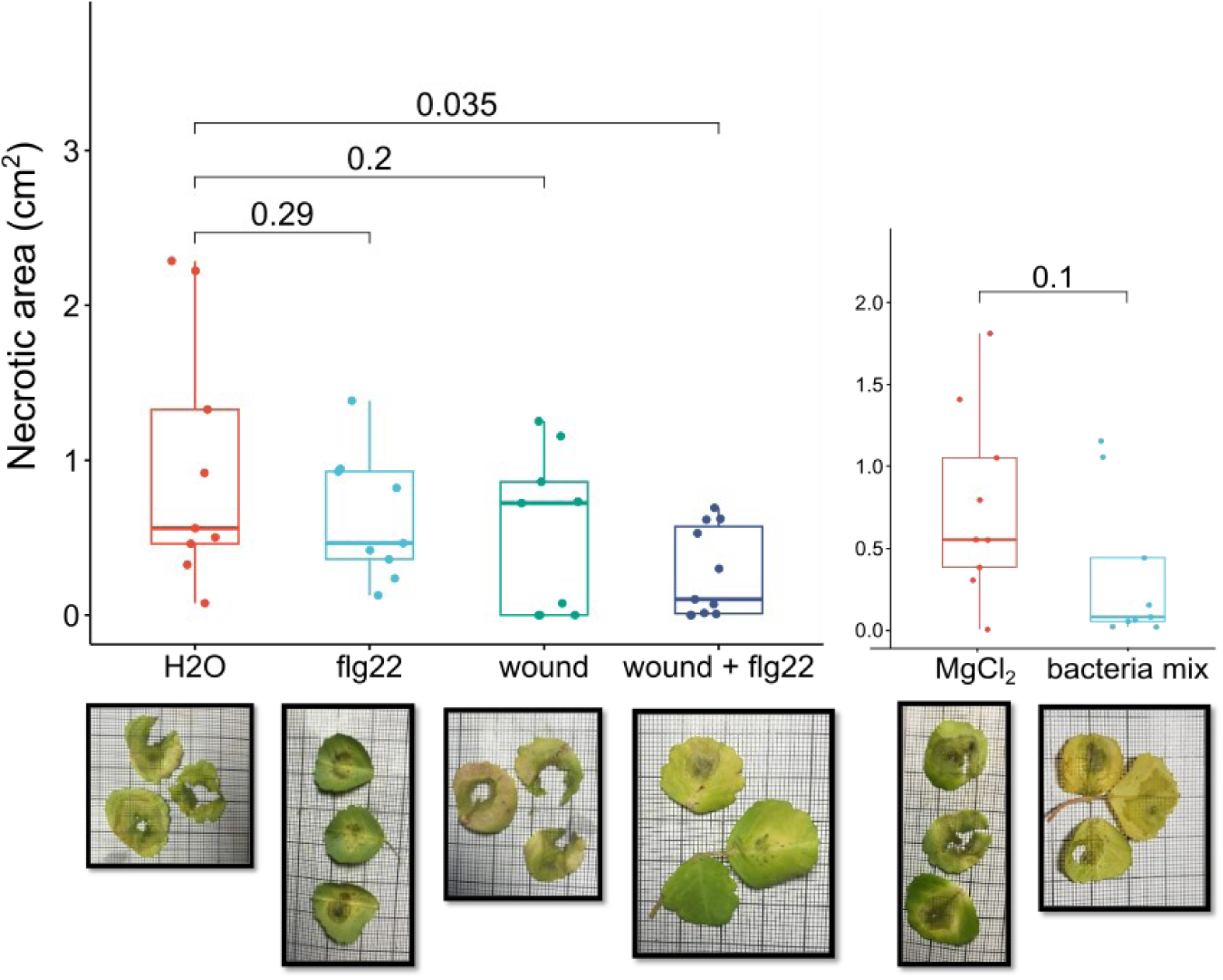
Size of *Botrytis cinerea* necrotic lesions (72hpi) on systemic leaves after stimulation of distal leaves with the following treatments: 5µL drop of H_2_O (control), 5 µL drop of 1µM flg22, wounding with forceps, wounding + 5 µL drop of 1µM flg22 as well as infiltrations10mM MgCl2 (control), live suspension of *P. syringae, D. chrysanthemi, E. rhapontici, P. agglomerans* at OD_660_=0.0005 each in 10mM MgCl_2_. All comparisons were performed at the same time. Box-plots depict distribution of individual data-points and means. ANOVA (F=2,86; p=0.18) suggest significant differences among the treatments and p-values from the Dunnet’s two tailed t-test pairwise comparisons are depicted on top of the plots.

**Supplemental Figure 3.**
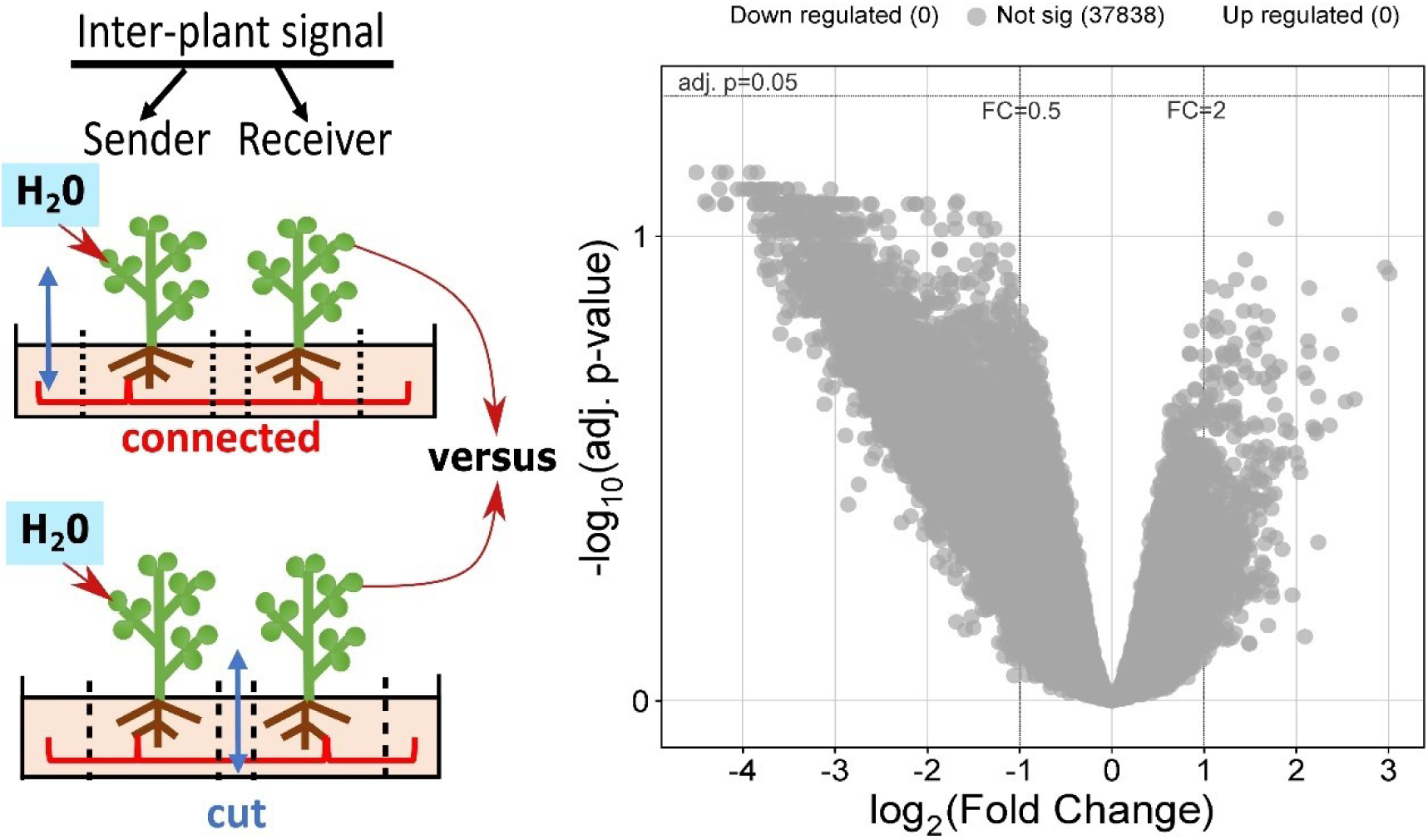
No significantly differentially expressed genes (DEGs) were identified when comparing responder plants from CNM connected plant and cut-CNM pots after sender plant treatment with water only. Volcano plot displays log_10_ of p-adj. and log_2_ of fold change for each transcript in the comparison.

**Supplemental Figure 4.**
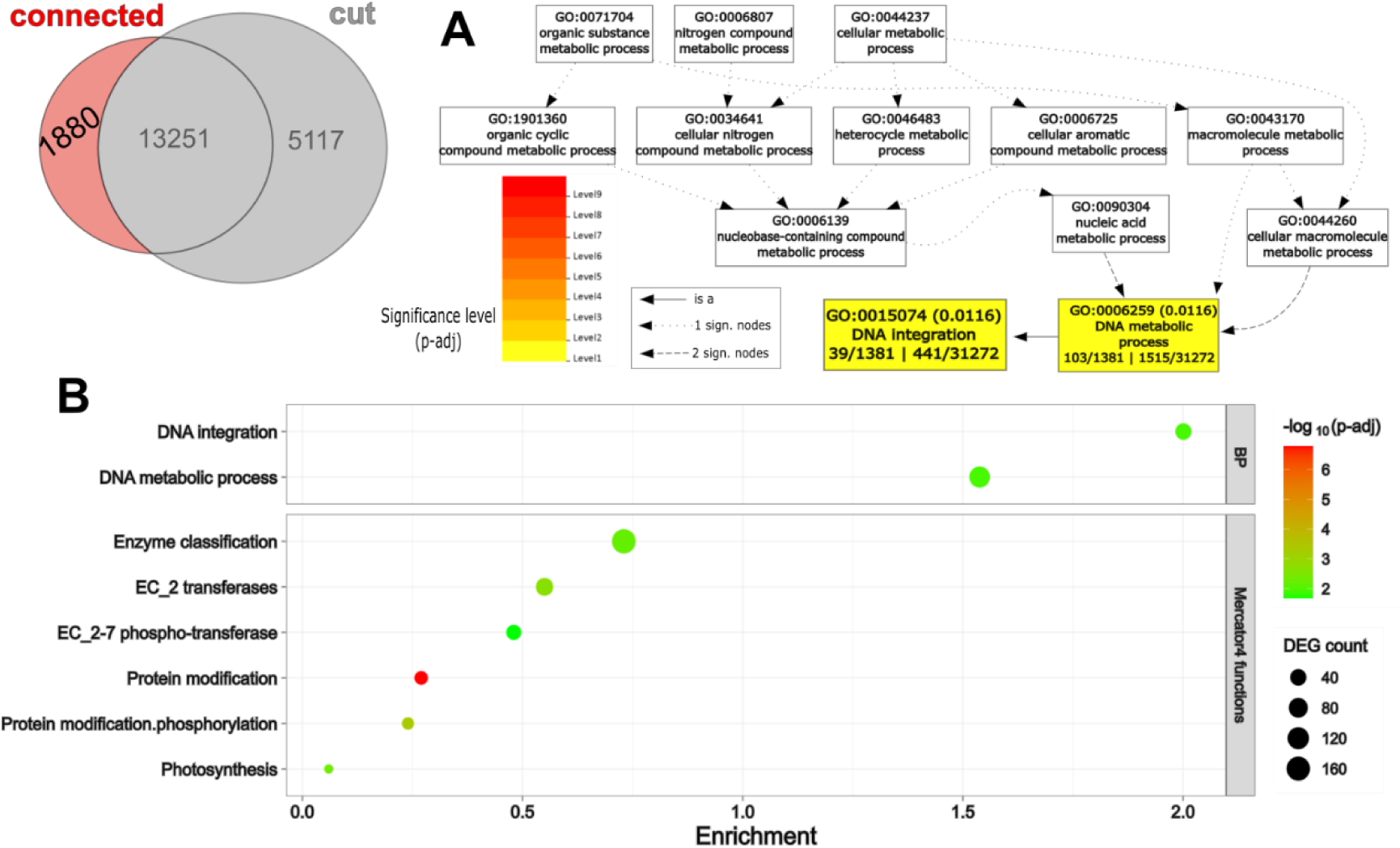
Functional enrichment of GO-terms with 1880 upregulated responder plant DEGs in CMN connected plant pots. (**A**) GO-identifiers and their known or inferred hierarchical interactions. Boxes contain colour-coded FDR-corrected p-values, the number DEGs within the GO-term from the query list/total GO-annotations in the query list | total number genes corresponding to the GO-term/ total GO-annotations in the genome. (**B**) Bubble plots depicting the enrichment factor, number of DEGs and log10 normalized FDR-corrected (adj.) p-value for the enrichment significance. Only GO biological functions (BP) and Mercator4 functional predictions with p-adj.<0.05 are displayed. Full list of GO-terms and bins for functional enrichment analysis is available in **Supplemental Table 2 tab-A** (for BP) and **tab D** (for Mercator4 annotation). The BP enrichment is based on the available functional annotations for 73.6% or 1384 out of the 1880 upregulated DEGs (**Supplemental Table 2 tab-C**).

**Supplemental Figure 5.**
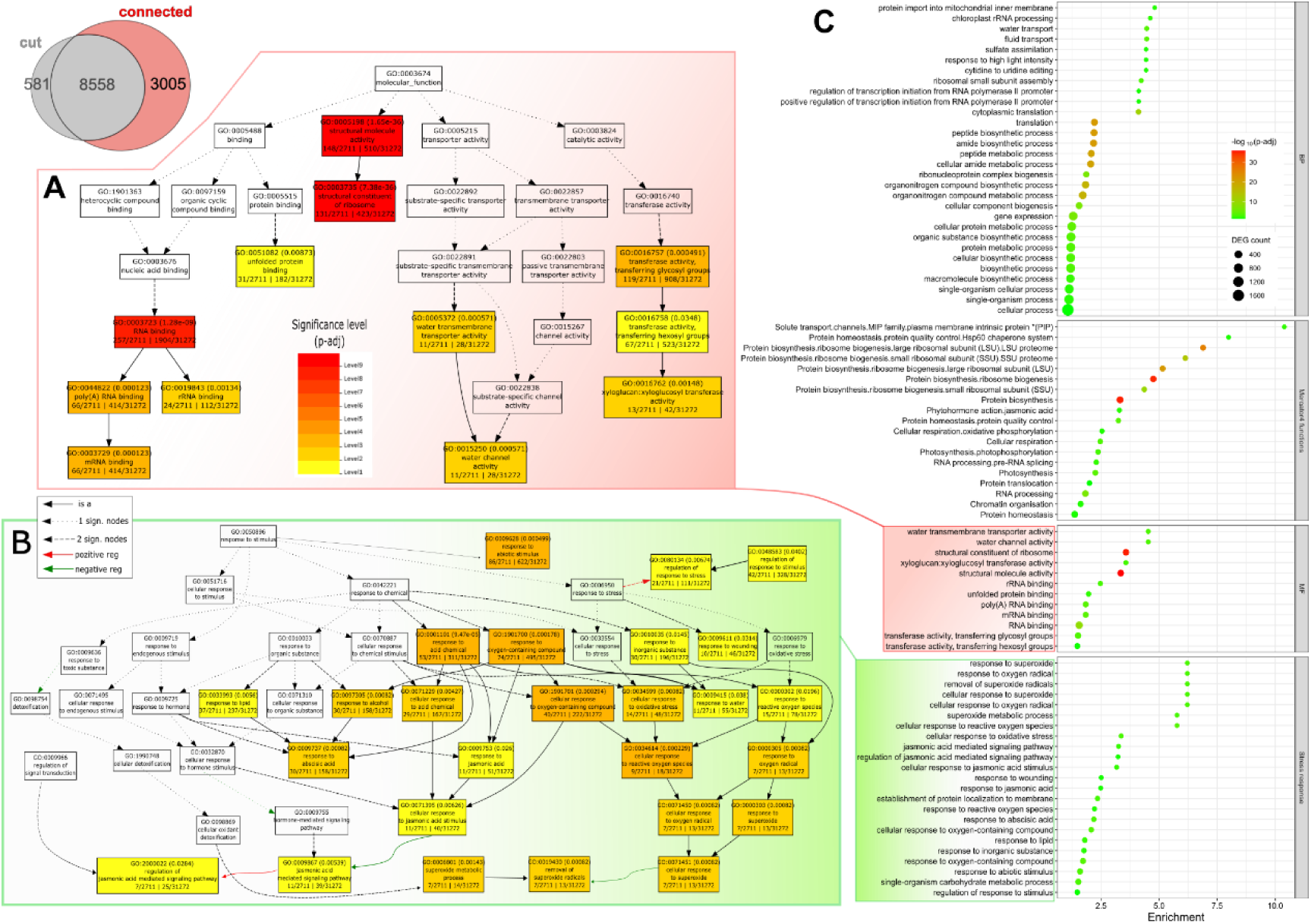
Functional enrichment of GO-terms with 3005 downregulated responder plant DEGs in CMN connected plant pots. (**A-B**) GO-identifiers and their known or inferred hierarchical interactions. Boxes contain colour-coded FDR-corrected p-values, the number DEGs within the GO-term from the query list/total GO-annotations in the query list | total number genes corresponding to the GO-term/ total GO-annotations in the genome. Different arrow types indicate GO-terms associated to 1 or 2 other significant enriched functions as well as known positive or negative regulation between genes within the associated GO-term. (**A**) Enriched GO-identifiers for molecular functions (MF) related to RNA binding and ribosome functions. (**B**) Enriched GO-identifiers for biological processes related to ROS metabolism, response to stimuli and jasmonic acid, collectively displayed under Stress response cluster. (**C**) Bubble plots depicting the enrichment factor, number of DEGs and log10 normalized FDR-corrected (adj.) p-value for the enrichment significance. For brevity the plot displays the top 30 most sign. Enriched GO biological functions (BP subpanel). All GO-terms with p-adj.<0.05 are displayed in the Stress response, MF and Mercator4 subpanels. Full list of GO-term descriptor for enrichment analysis is available in **Supplemental Table 2 tab-B** (for BP and MF) and **tab E** (for Mercator4 annotation). The displayed **Stress response** category represents a subset of BP GO-terms that were manually assigned to this category based on the description semantics and are fully listed in **Supplemental Table 2 tab-B.** The BP and MF enrichment is based on the available functional annotations for 90.3% or 2713 out of the 3005 DEGs (**Supplemental Table 2 tab-C**).

**Supplemental Figure 6.**
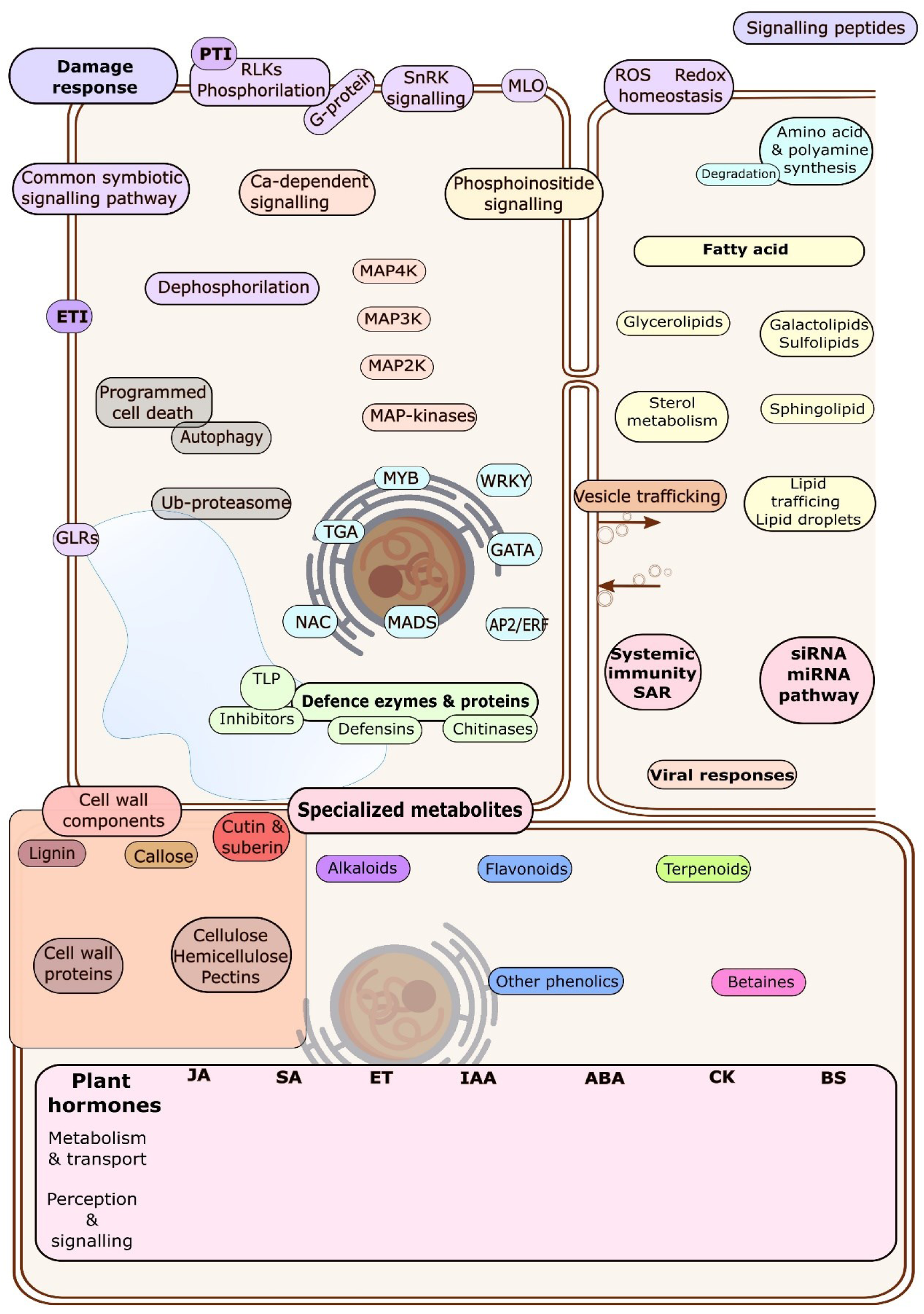
Image file for manually curated “Defence signalling & response” category to display fold change of genes corresponding to bins listed in **Supplemental Table 3 tab-C.**

**Supplemental Figure 7.**
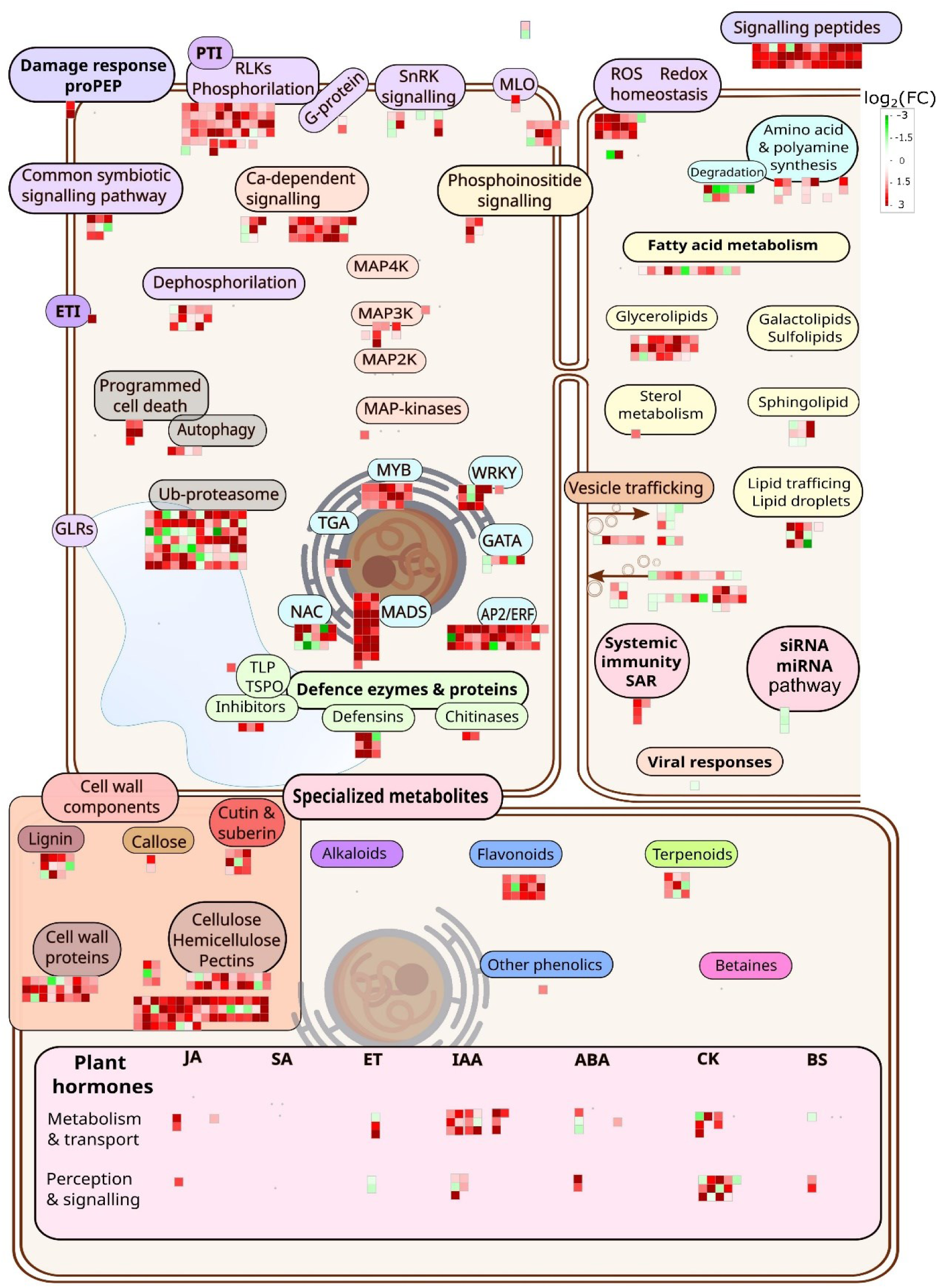
Plant “Defence signalling & response” pathways genes are upregulated in receivers with cut CMN. Log2 (transcript fold) changes of plant functions encoded by up- (red) and down-regulated (green) DEGs in receivers of wound and flg22 signals compared to H_2_O controls.

**Supplemental Figure 8.**
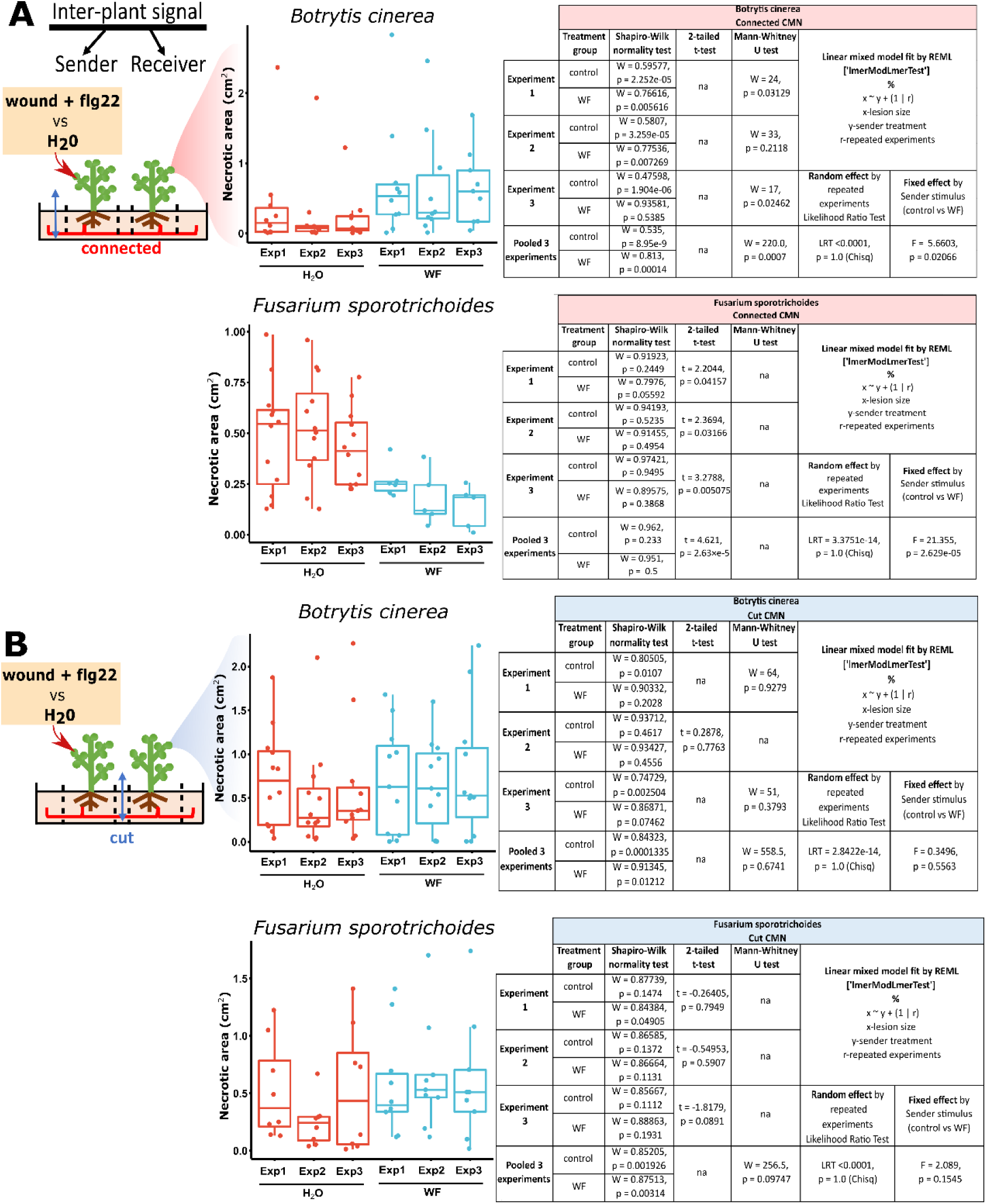
Size of necrotic lesions on *Mediago truncatula* receiver leaves 72h after inoculation with *Botrytis cinerea* and *Fusarium sporotichoides* in CMN-connected (A) and cut CMN treatments (B). Experiments in panel A and B were performed independently three times, each with 10-11 biological replicates. Line indicates the median, boxes represent the interquartile range (IQR), whiskers - variance within 1.5xIQR. Table displays test-statistics for each individual experiment as well as the pooled data represented in the **Figure 4**. Mann-Whitney U-test was performed where data deviate from normal distribution after Shapiro-Wilk test. Two-tailed t-test was performed on normally distributed data. Pooled data were analysed using linear mixed model (lmer). Results did not differ among repeated experiments for any of the treatments as demonstrated by Likelihood Ratio Test (LRT) and non-significant p-values for the associated Chi-square test. Only sender plant treatment displayed significant effect on receiver lesion size in the connected CMN treatment (**A**).

**Supplemental Figure 9.**
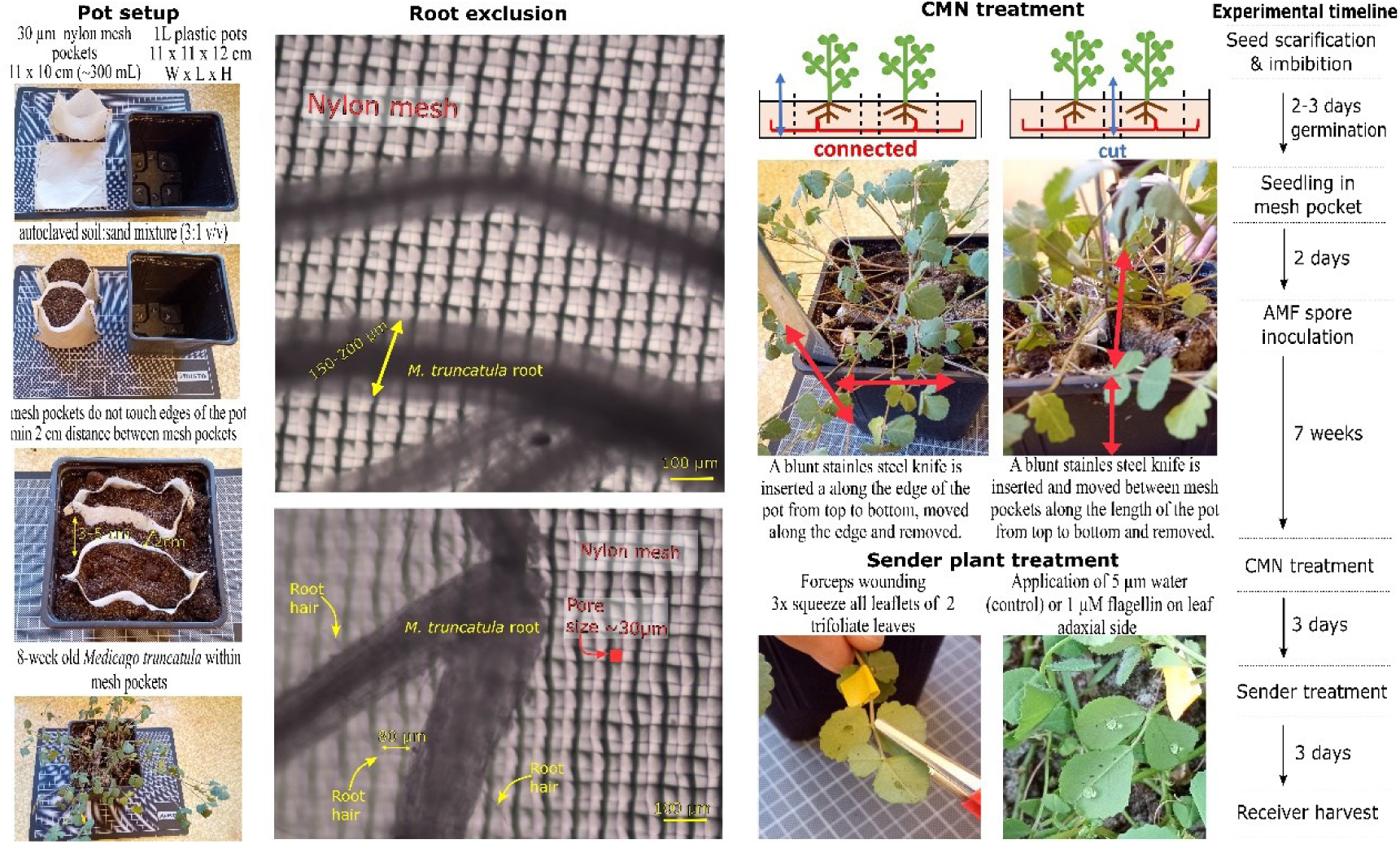
Visual representation of the experimental setup and treatments. The distance between the mesh pockets is >2cm in the middle of the pots and increases towards the edges (3-5cm) due to the geometry of soil-filled mesh pockets. *Medicago truncatula* fine roots are 150-200 µm in diameter and cannot penetrate through the 30 µm mesh or be injured during the knife treatment of CMN. The root hairs may penetrate through the pore size but due to their short length (50-80 µm) and the distance between the plants (>2cm) they cannot make physical inter-plant contact.

**Supplemental figure 10.**
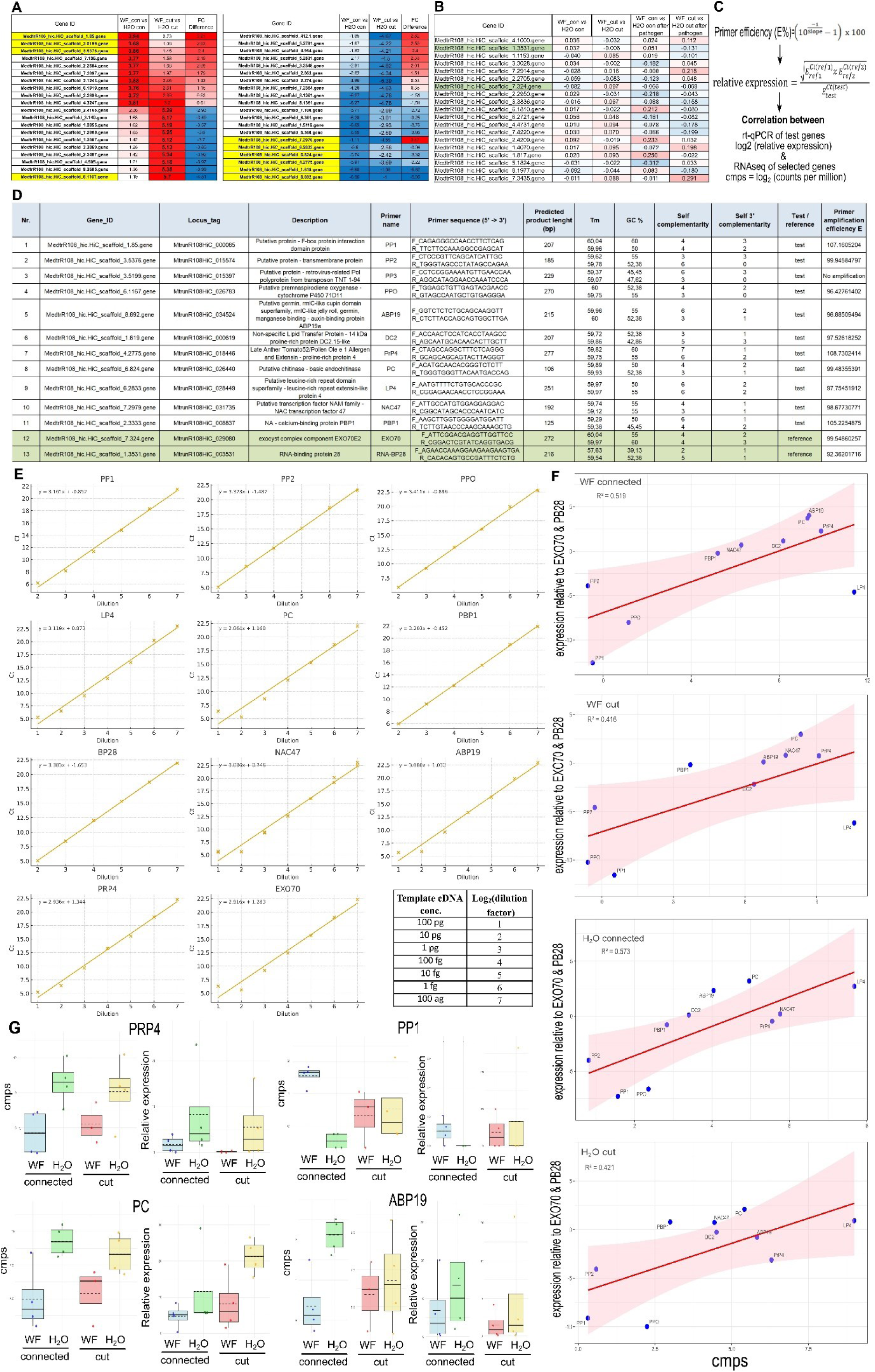
Correlation of gene expression data 1126 from RNA-seq experiment and an independent sample set. **(A)** Test genes were selected based on the highest log2FC in receivers from wound and flg22 (WF) treated plants with intact or cut CMN as well as the largest relative differences between the “connected” and “cut” treatments. **(B)** Candidate reference genes were selected among the genes with the lowest FC and no sign. FC among any of the treatments. Selected references were EXO70 and RNA-BP28. **(C)** Analysis pipeline and equations used for calculating the primer efficiency and relative expression. **(D)** Full list of the selected test and reference gene primers and gene descriptors. Primer efficiency was first calculated using a dilution series of gel-purified cDNA amplicons displayed in **(E)**. Relative expression was calculated as the geometric mean of the amplification of two reference genes divided by the amplification of a test gene **(C)**. The correlation between the relative expression of selected targets from independent samples by rt-qPCR and RNA-seq is depicted in panel **(F)**. The expression pattern of representative set of test genes in the “connected” and “cut” systems after WF and H_2_O treatments is similar in RNA-seq and qPCR analysis from independent samples. CMPS (plural) refers to counts per million from *limma* package output files.

**Supplemental figure 11.**
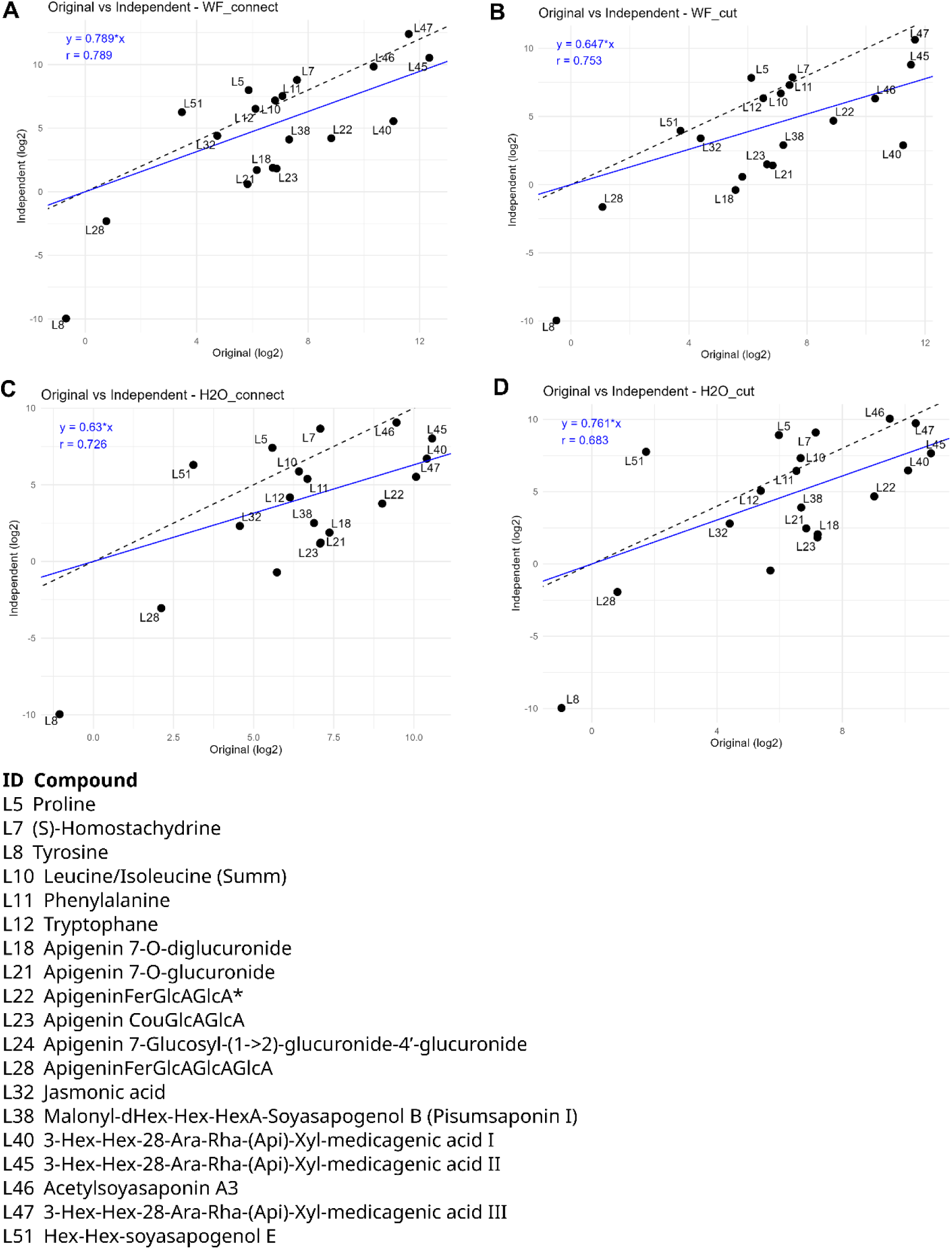
Correlation of metabolite levels from two independent repeated experiments. The “original” sample data are presented in Figure 3. The “independent” samples were obtained from an independent repeated experiment. Comparisons between the “original” and “independent” samples include intact **(A)** or cut **(B)** CMN between receivers and wound and flg22 (WF) treated senders as well as intact **(C)** or cut **(D)** CMN between receivers and water-treated senders. Dashed black line corresponds to perfect correlation between “original” and “independent” samples. Blue line and the equation indicate fitted regression line for the correspondence of both datasets along with Pearson correlation (r). The compound quantities (μg/g FW) were determined using the available standards and available in **Supplemental Table 4 tab-C**.

## Literature Cited

1. Orlovskis, Z. & Reymond, P. Pieris brassicae eggs trigger interplant systemic acquired resistance against a foliar pathogen in Arabidopsis. New Phytol. 228, 1652–1661 (2020).

2. Riedlmeier, M. et al. Monoterpenes Support Systemic Acquired Resistance within and between Plants. Plant Cell 29, 1440–1459 (2017).

3. Bais, H. P., Park, S.-W., Weir, T. L., Callaway, R. M. & Vivanco, J. M. How plants communicate using the underground information superhighway. Trends Plant Sci. 9, 26–32 (2004).

4. Barto, E. K., Weidenhamer, J. D., Cipollini, D. & Rillig, M. C. Fungal superhighways: do common mycorrhizal networks enhance below ground communication? Trends Plant Sci. 17, 633–637 (2012).

5. Brundrett, M. C. & Tedersoo, L. Evolutionary history of mycorrhizal symbioses and global host plant diversity. New Phytol. 220, 1108–1115 (2018).

6. Read, D. J. et al. Photosynthate transfer from an autotrophic orchid to conspecific heterotrophic protocorms through a common mycorrhizal network. New Phytol. 243, 398–406 (2024).

7. Francis, R. & Read, D. J. Direct transfer of carbon between plants connected by vesicular–arbuscular mycorrhizal mycelium. Nature 307, 53–56 (1984).

8. Simard, S. W. et al. Net transfer of carbon between ectomycorrhizal tree species in the field. Nature 388, 579–582 (1997).

9. Song, Y. Y. et al. Interplant communication of tomato plants through underground common mycorrhizal networks. PloS One 5, e13324 (2010).

10. Babikova, Z. et al. Underground signals carried through common mycelial networks warn neighbouring plants of aphid attack. Ecol. Lett. 16, 835–843 (2013).

11. Pieterse, C. M. J., Leon-Reyes, A., Van der Ent, S. & Van Wees, S. C. M. Networking by small-molecule hormones in plant immunity. Nat. Chem. Biol. 5, 308–316 (2009).

12. Zhou, F. et al. Co-incidence of Damage and Microbial Patterns Controls Localized Immune Responses in Roots. Cell 180, 440–453.e18 (2020).

13. Orlovskis, Z., Singh, A., Kliot, A., Huang, W. & Hogenhout, S. A. The phytoplasma SAP54 effector acts as a molecular matchmaker for leafhopper vectors by targeting plant MADS-box factor SVP. eLife 13, RP98992 (2025).

14. Khashi u Rahman, M., Zhou, X. & Wu, F. The role of root exudates, CMNs, and VOCs in plant–plant interaction. J. Plant Interact. 14, 630–636 (2019).

15. Song, Y. Y., Simard, S. W., Carroll, A., Mohn, W. W. & Zeng, R. S. Defoliation of interior Douglas-fir elicits carbon transfer and stress signalling to ponderosa pine neighbors through ectomycorrhizal networks. Sci. Rep. 5, 8495 (2015).

16. Rodriguez-Morelos, V. H., Calonne-Salmon, M., Bremhorst, V., Garcés-Ruiz, M. & Declerck, S. Fungicides With Contrasting Mode of Action Differentially Affect Hyphal Healing Mechanism in Gigaspora sp. and Rhizophagus irregularis. Front. Plant Sci. 12, 642094 (2021).

17. Richter, F., Calonne-Salmon, M., Heijden, M. G. A. van der, Declerck, S. & Stanley, C. E. AMF-SporeChip provides new insights into arbuscular mycorrhizal fungal asymbiotic hyphal growth dynamics at the cellular level. Lab. Chip 24, 1930–1946 (2024).

18. Schütz, L., Saharan, K., Mäder, P., Boller, T. & Mathimaran, N. Rate of hyphal spread of arbuscular mycorrhizal fungi from pigeon pea to finger millet and their contribution to plant growth and nutrient uptake in experimental microcosms. Appl. Soil Ecol. 169, 104156 (2022).

19. Szakiel, A., Pączkowski, C. & Henry, M. Influence of environmental biotic factors on the content of saponins in plants. Phytochem. Rev. 10, 493–502 (2011).

20. Da Silva, P. et al. High toxicity and specificity of the saponin 3-GlcA-28-AraRhaxyl-medicagenate, from Medicago truncatula seeds, for Sitophilus oryzae. BMC Chem. Biol. 12, 1–9 (2012).

21. Suzuki, H. et al. Lotus japonicus Triterpenoid Profile and Characterization of the CYP716A51 and LjCYP93E1 Genes Involved in Their Biosynthesis In Planta. Plant Cell Physiol. 60, 2496–2509 (2019).

22. D’Addabbo, T. et al. Activity of Saponins from Medicago Species against Phytoparasitic Nematodes. Plants 9, 443 (2020).

23. Norvienyeku, J. et al. Bayogenin 3-O-cellobioside confers non-cultivar-specific defence against the rice blast fungus Pyricularia oryzae. Plant Biotechnol. J. 19, 589–601 (2021).

24. Yan, D.-H. et al. Antifungal Activities of Volatile Secondary Metabolites of Four Diaporthe Strains Isolated from Catharanthus roseus. J. Fungi 4, 65 (2018).

25. Spaepen, S., Vanderleyden, J. & Remans, R. Indole-3-acetic acid in microbial and microorganism-plant signaling. FEMS Microbiol. Rev. 31, 425–448 (2007).

26. McClerklin, S. A. et al. Indole-3-acetaldehyde dehydrogenase-dependent auxin synthesis contributes to virulence of Pseudomonas syringae strain DC3000. PLOS Pathog. 14, e1006811 (2018).

27. Ranocha, P. et al. Arabidopsis WAT1 is a vacuolar auxin transport facilitator required for auxin homoeostasis. Nat. Commun. 4, 2625 (2013).

28. Fukushima, E. O. et al. Combinatorial Biosynthesis of Legume Natural and Rare Triterpenoids in Engineered Yeast. Plant Cell Physiol. 54, 740–749 (2013).

29. Qamar, A., Mysore, K. & Senthil-Kumar, M. Role of proline and pyrroline-5-carboxylate metabolism in plant defense against invading pathogens. Front. Plant Sci. 6, (2015).

30. Siddique, A., Kandpal, G. & Kumar, P. Proline Accumulation and its Defensive Role Under Diverse Stress Condition in Plants: An Overview. J. Pure Appl. Microbiol. 12, 1655–1659 (2018).

31. Pélissier, R. et al. Plant neighbour-modulated susceptibility to pathogens in intraspecific mixtures. J. Exp. Bot. 72, 6570–6580 (2021).

32. Alfonso, E. et al. Insect eggs trigger systemic acquired resistance against a fungal and an oomycete pathogen. New Phytol. 232, 2491–2505 (2021).

33. Erb, M. & Reymond, P. Molecular Interactions Between Plants and Insect Herbivores. Annu. Rev. Plant Biol. 70, 527–557 (2019).

34. Chisholm, S. T., Coaker, G., Day, B. & Staskawicz, B. J. Host-Microbe Interactions: Shaping the Evolution of the Plant Immune Response. Cell 124, 803–814 (2006).

35. Wenig, M. et al. Systemic acquired resistance networks amplify airborne defense cues. Nat. Commun. 10, 3813 (2019).

36. Jia, Q., et al. Origin and early evolution of the plant terpene synthase family. Proc. Natl. Acad. Sci. 119, e2100361119 (2022).

37. Jiang, S.-Y., Jin, J., Sarojam, R. & Ramachandran, S. A Comprehensive Survey on the Terpene Synthase Gene Family Provides New Insight into Its Evolutionary Patterns. Genome Biol. Evol. 11, 2078–2098 (2019).

38. Wang, Q. et al. Expansion and functional divergence of terpene synthase genes in angiosperms: a driving force of terpene diversity. Hortic. Res. 12, uhae272 (2025).

39. St-Arnaud, M., Hamel, C., Vimard, B., Caron, M. & Fortin, J. A. Enhanced hyphal growth and spore production of the arbuscular mycorrhizal fungus Glomus intraradices in an in vitro system in the absence of host roots. Mycol. Res. 100, 328–332 (1996).

40. Baranski, R. Genetic Transformation of Carrot (Daucus carota) and Other Apiaceae Species. Transgenic Plant J. 2, 18–38 (2008).

41. Ron, M. et al. Hairy Root Transformation Using Agrobacterium rhizogenes as a Tool for Exploring Cell Type-Specific Gene Expression and Function Using Tomato as a Model. Plant Physiol. 166, 455–469 (2014).

42. Vierheilig, H., Coughlan, A. P., Wyss, U. & Piché, Y. Ink and Vinegar, a Simple Staining Technique for Arbuscular-Mycorrhizal Fungi. Appl. Environ. Microbiol. 64, 5004–5007 (1998).

43. Dobin, A. & Gingeras, T. R. Mapping RNA-seq Reads with STAR. Curr. Protoc. Bioinforma. Ed. Board Andreas Baxevanis Al 51, 11.14.1–11.14.19 (2015).

44. Kopylova, E., Noé, L. & Touzet, H. SortMeRNA: fast and accurate filtering of ribosomal RNAs in metatranscriptomic data. Bioinformatics 28, 3211–3217 (2012).

45. Ritchie, M. E. et al. limma powers differential expression analyses for RNA-sequencing and microarray studies. Nucleic Acids Res. 43, e47 (2015).

46. Tian, T. et al. agriGO v2.0: a GO analysis toolkit for the agricultural community, 2017 update. Nucleic Acids Res. 45, W122–W129 (2017).

47. Lohse, M. et al. Mercator: a fast and simple web server for genome scale functional annotation of plant sequence data: Mercator: sequence functional annotation server. Plant Cell Environ. 37, 1250–1258 (2014).

48. Thimm, O. et al. mapman: a user-driven tool to display genomics data sets onto diagrams of metabolic pathways and other biological processes. Plant J. 37, 914–939 (2004).

49. Usadel, B. et al. PageMan: an interactive ontology tool to generate, display, and annotate overview graphs for profiling experiments. BMC Bioinformatics 7, 535 (2006).

50. Usadel, B. et al. Extension of the Visualization Tool MapMan to Allow Statistical Analysis of Arrays, Display of Coresponding Genes, and Comparison with Known Responses. Plant Physiol. 138, 1195–1204 (2005).

