## Supplementary figures and images for "Common mycelial network modulates neighbour-primed plant defences against foliar pathogens by co-opting distinct inter-plant metabolic and biotic stress responses"

### Supplemental Figure 1

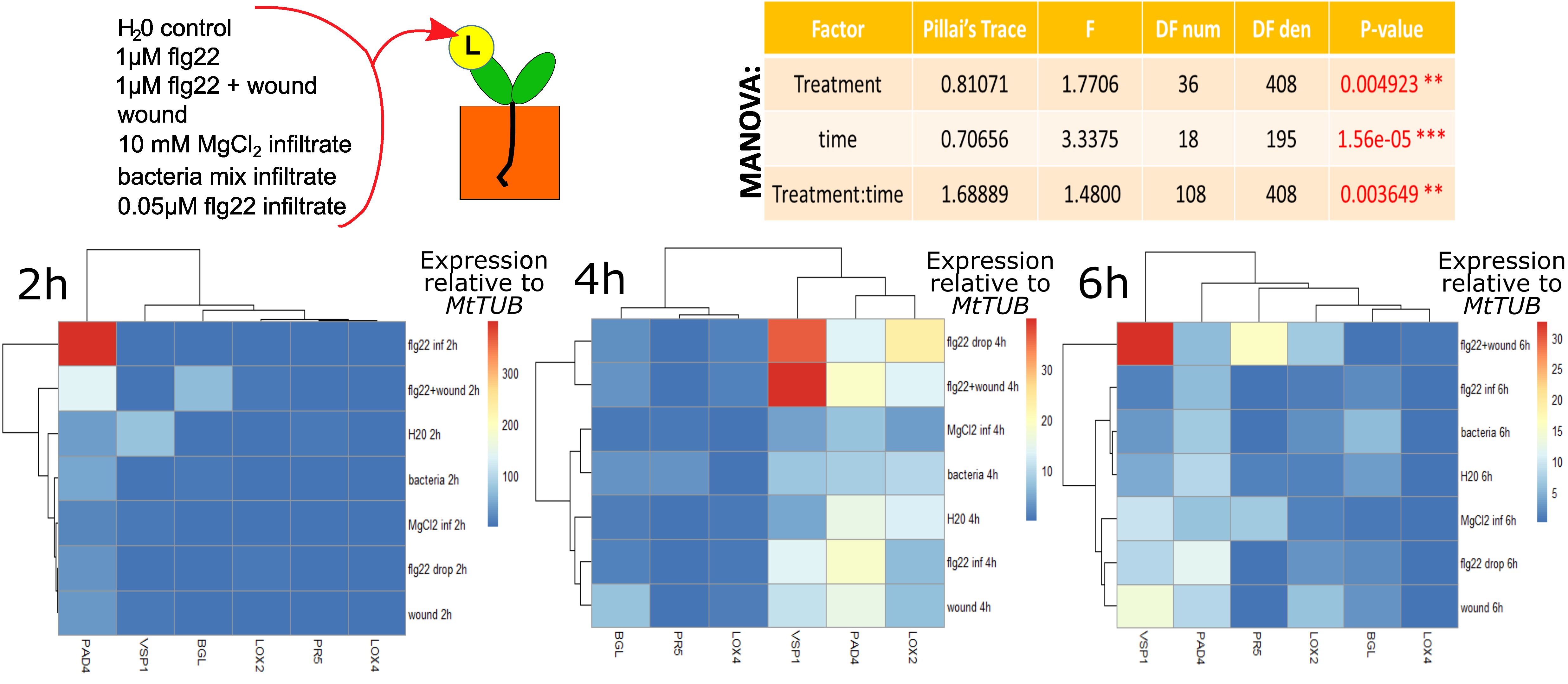

### Supplemental Figure 2

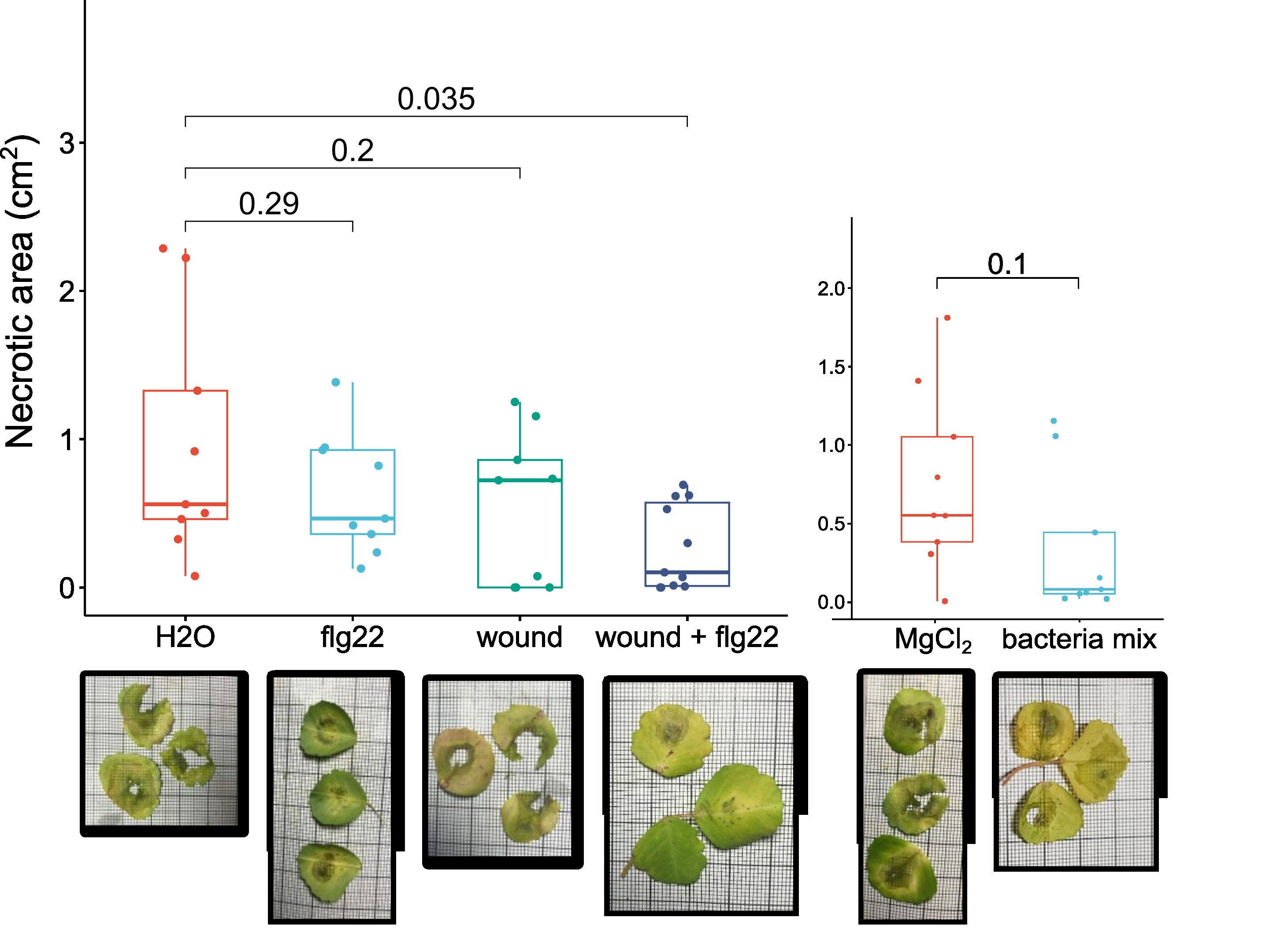

### Supplemental Figure 3

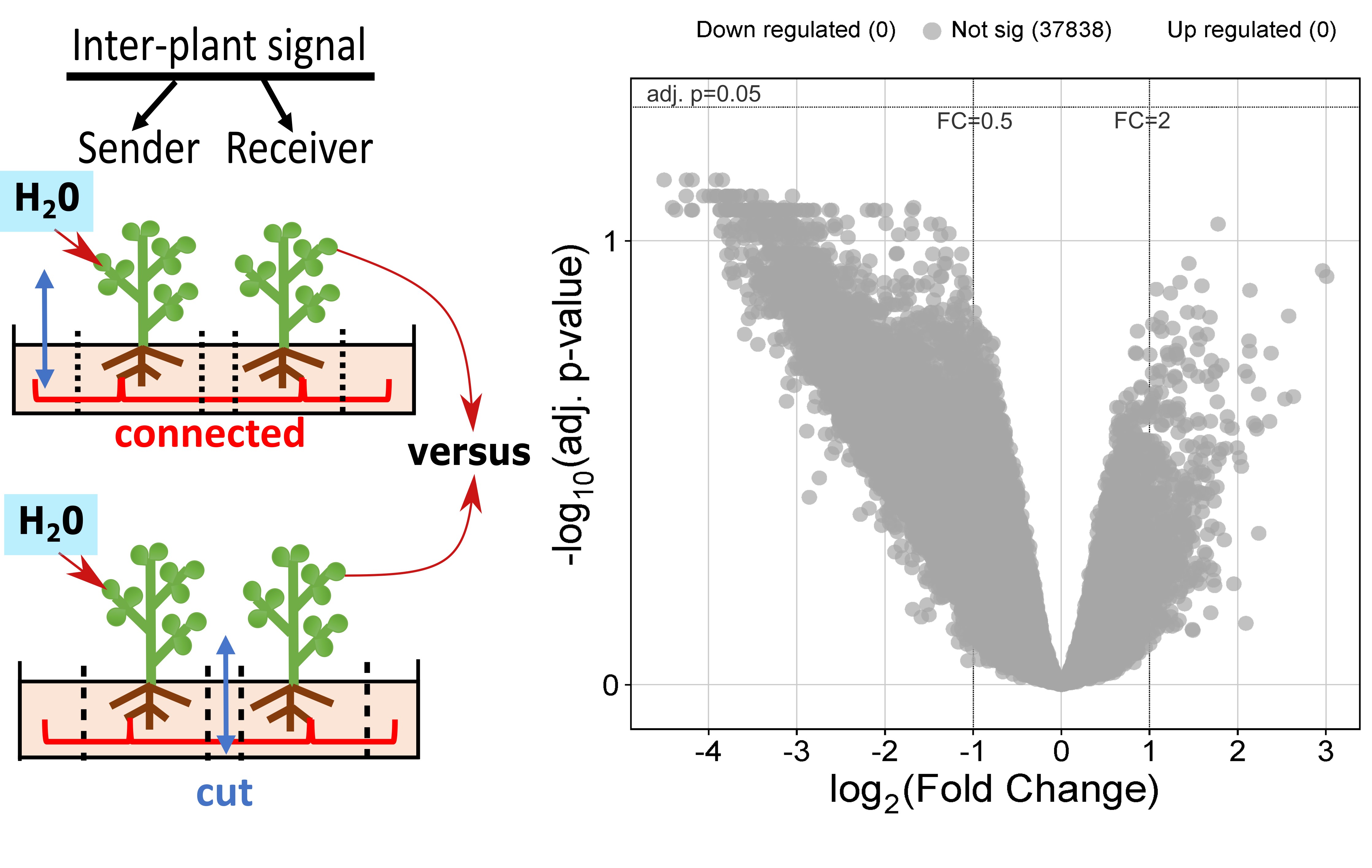

### Supplemental Figure 4

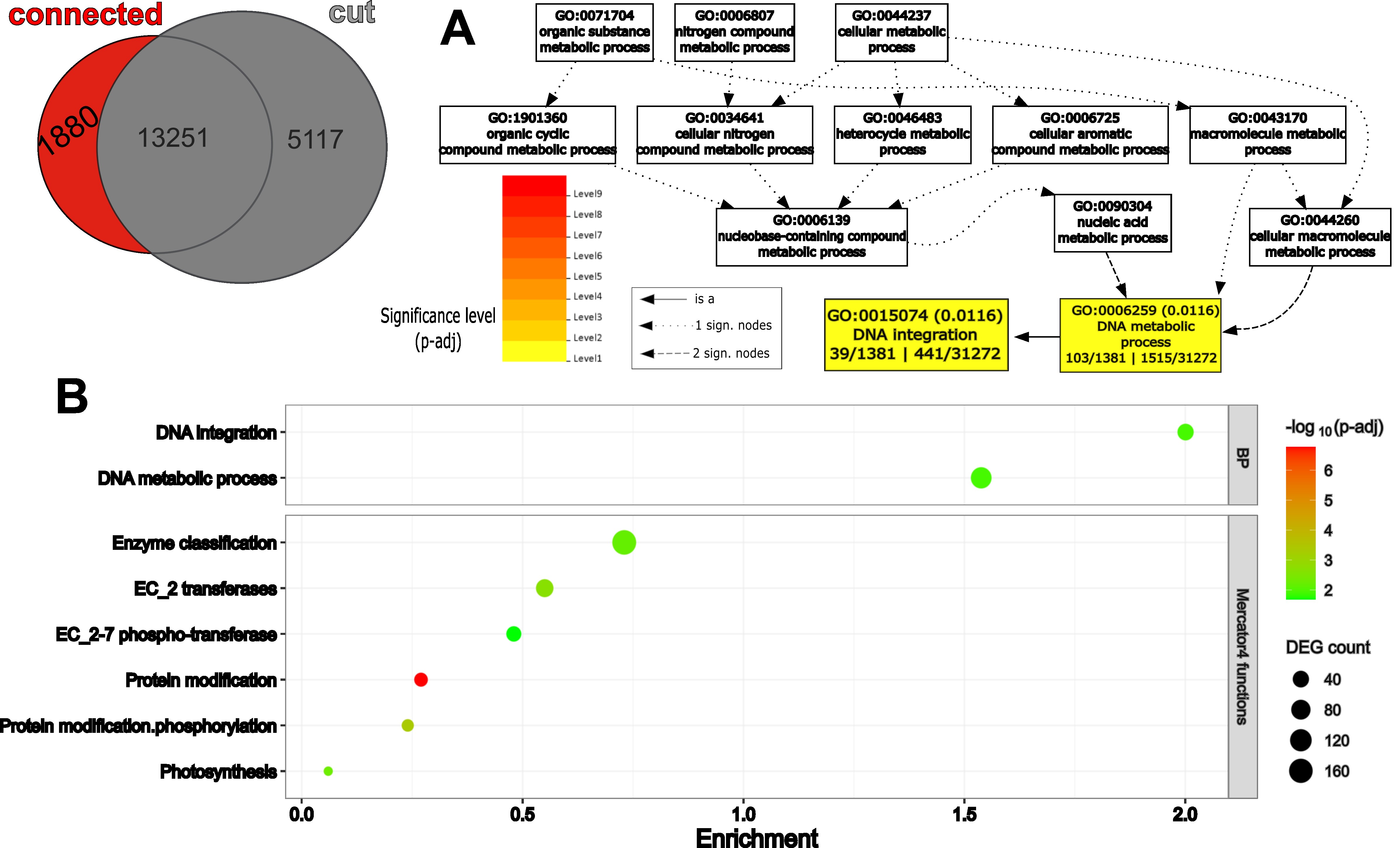

### Supplemental Figure 5

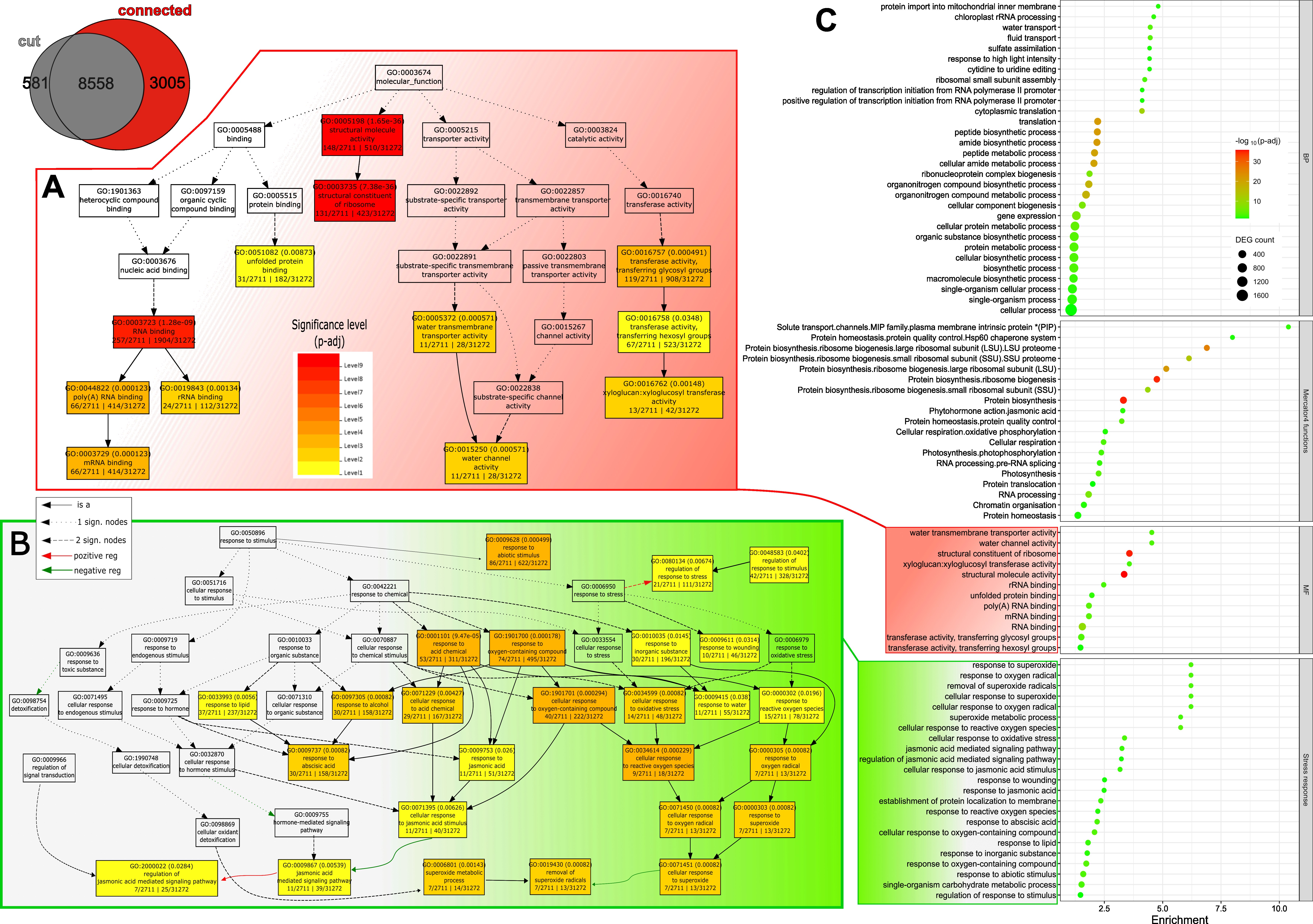

### Supplemental Figure 6

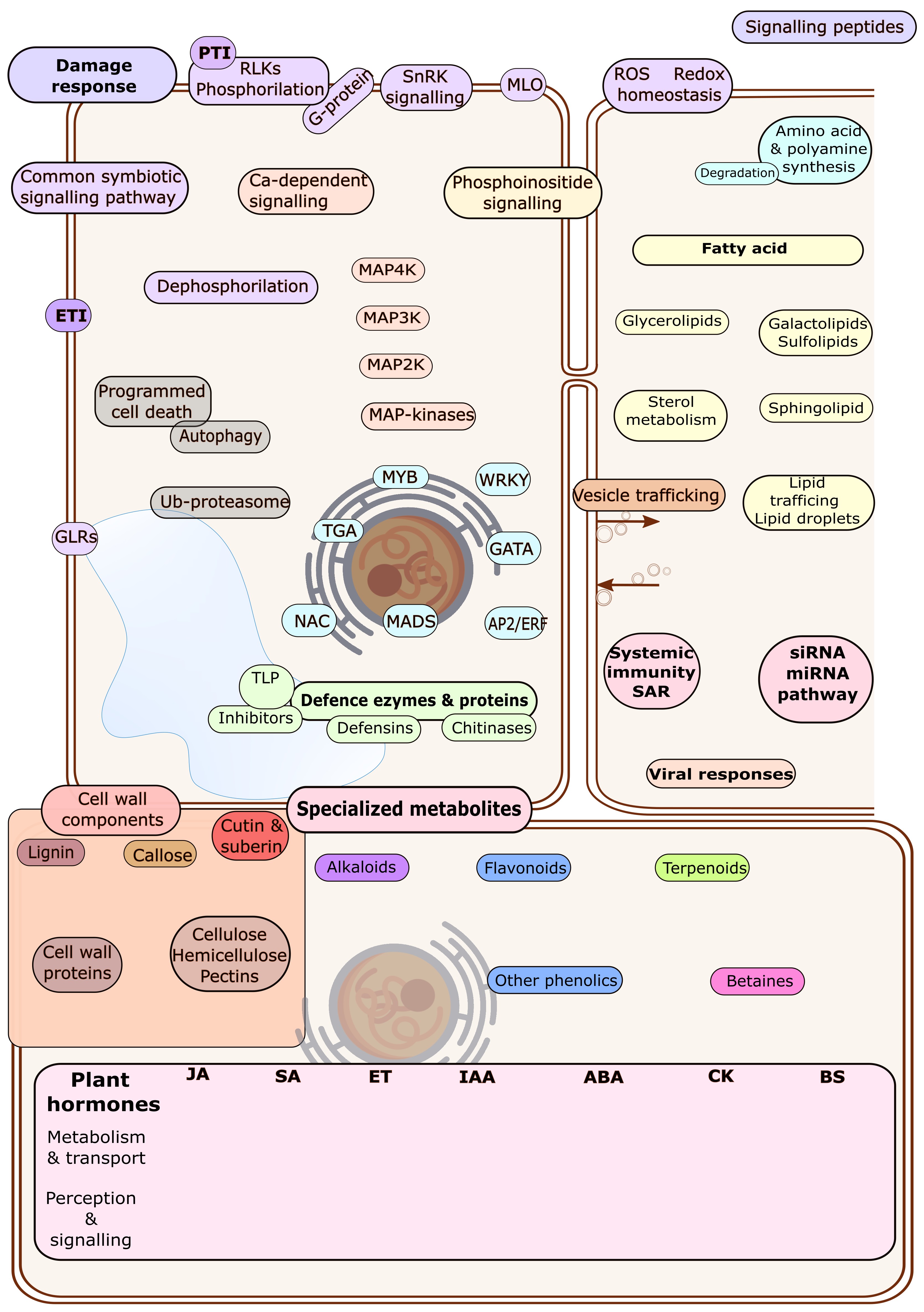

### Supplemental Figure 7

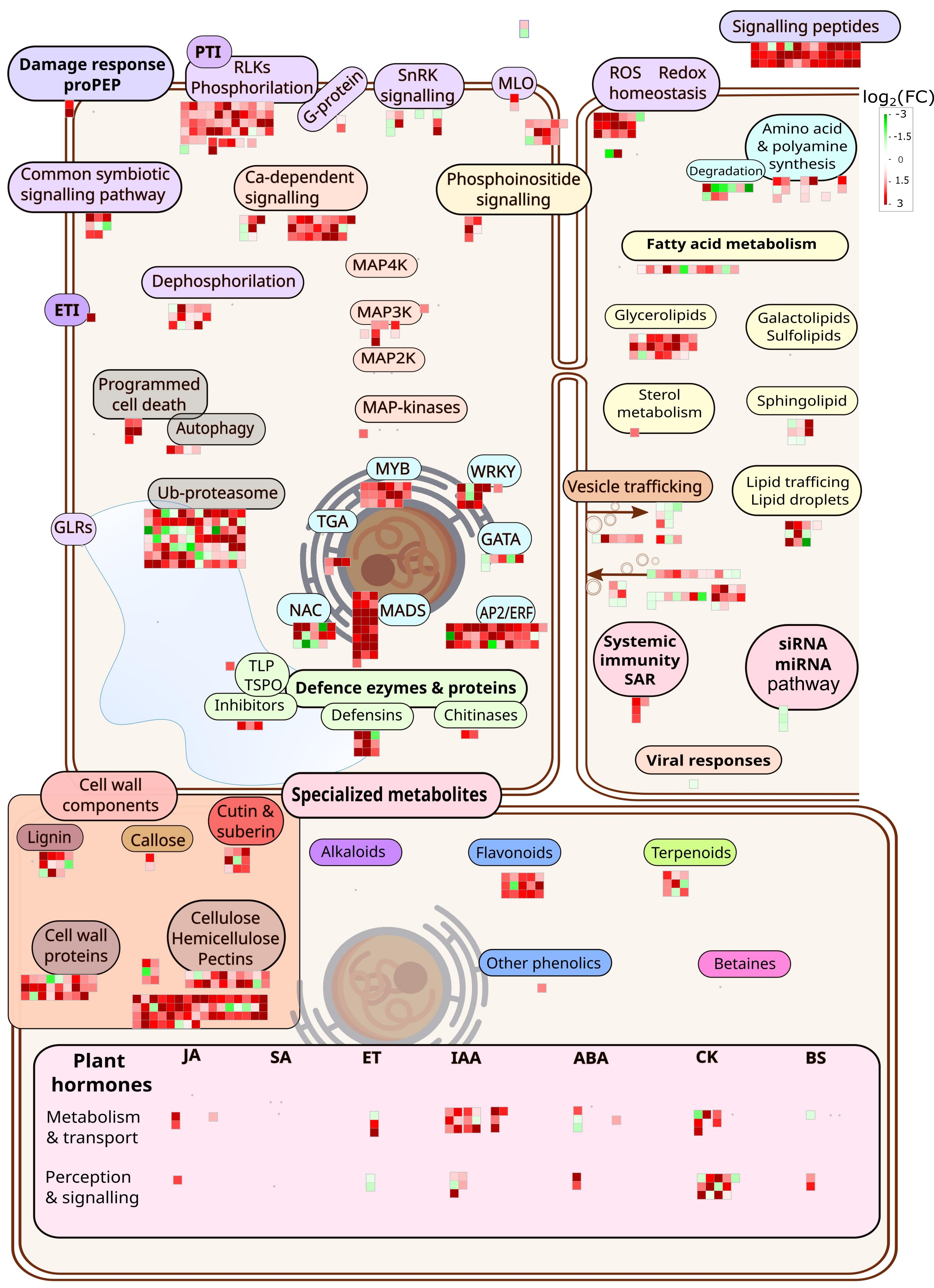

### Supplemental Figure 8

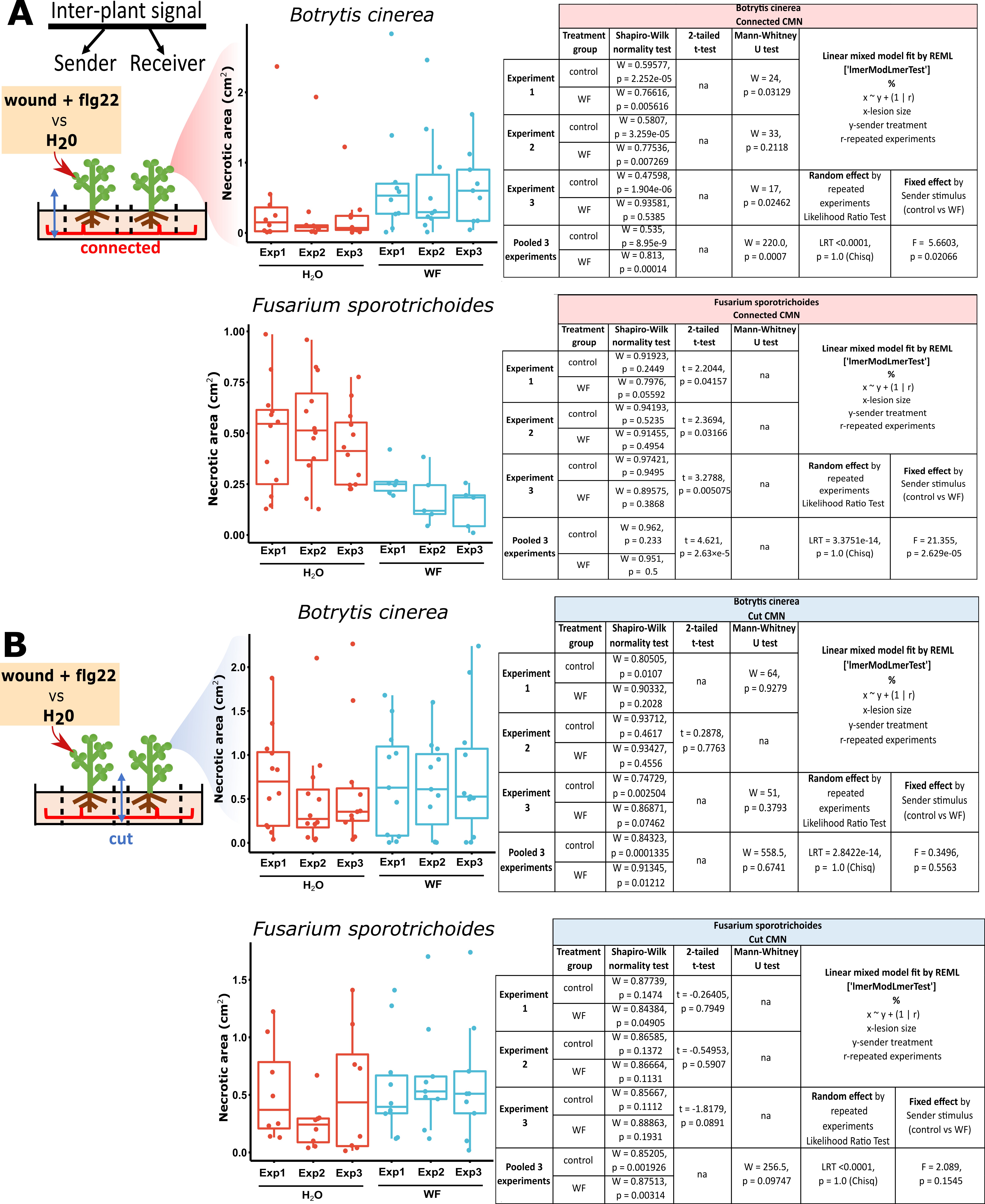

### Supplemental Figure 9

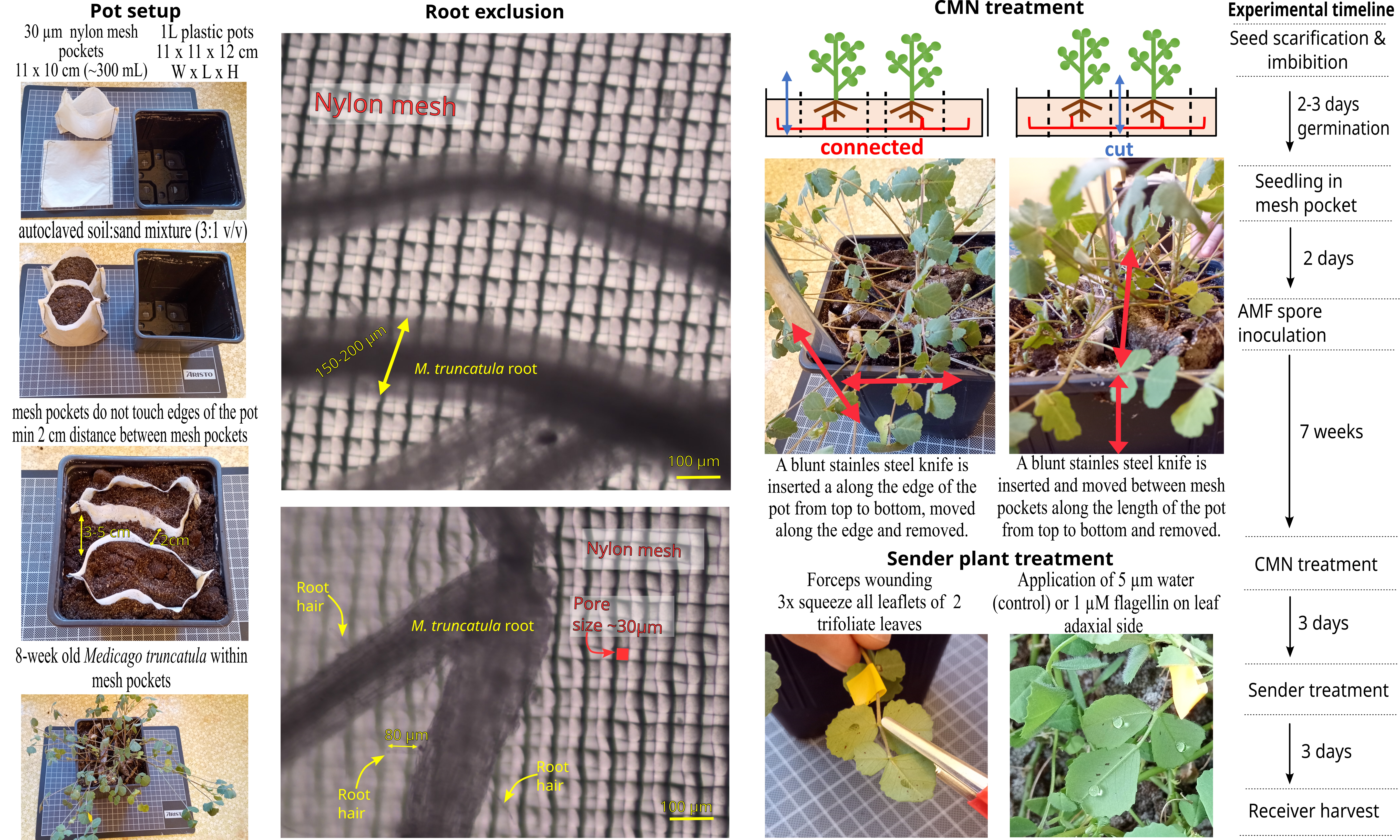

### Supplemental Figure 10

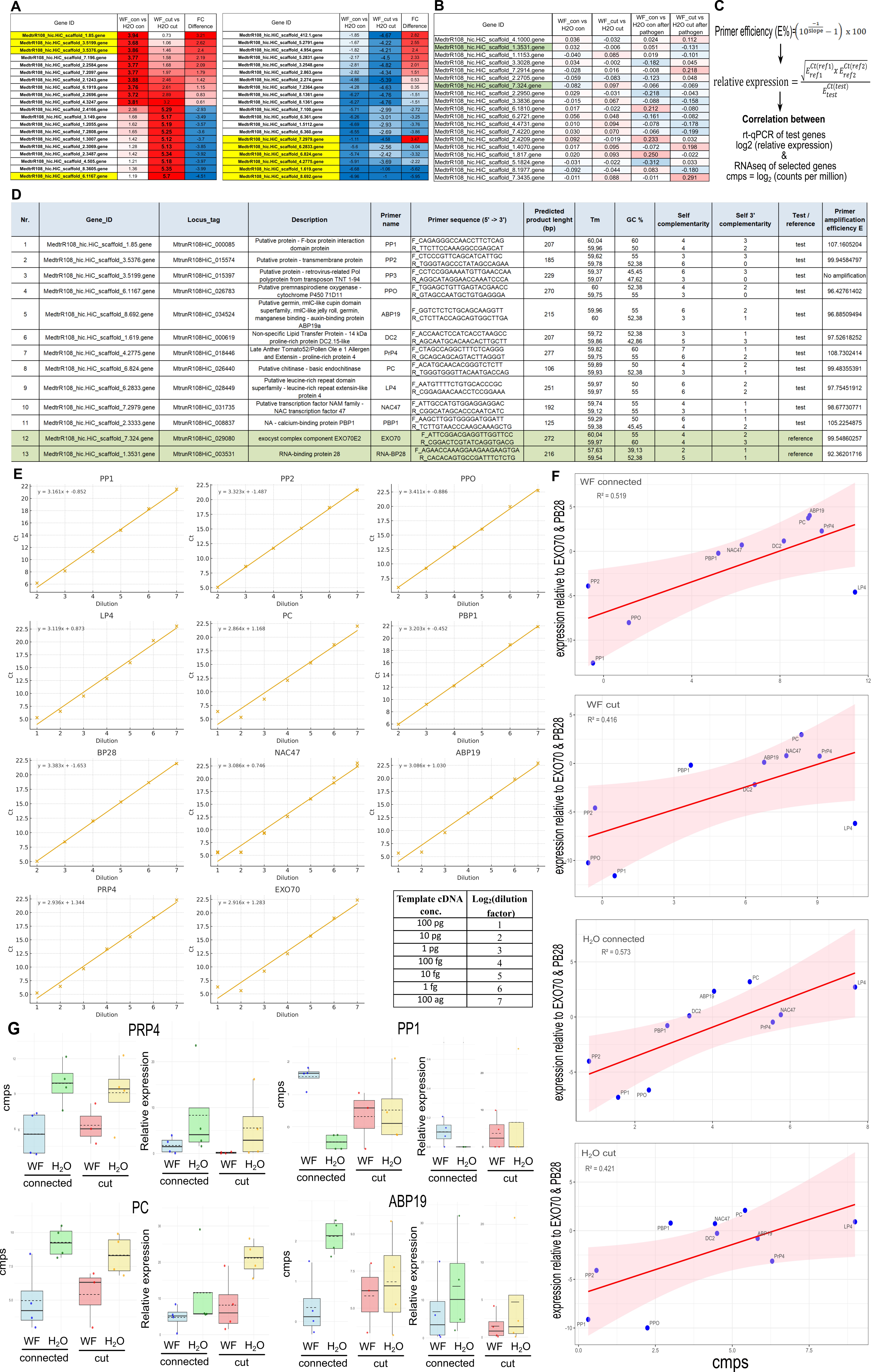

### Supplemental Figure 11

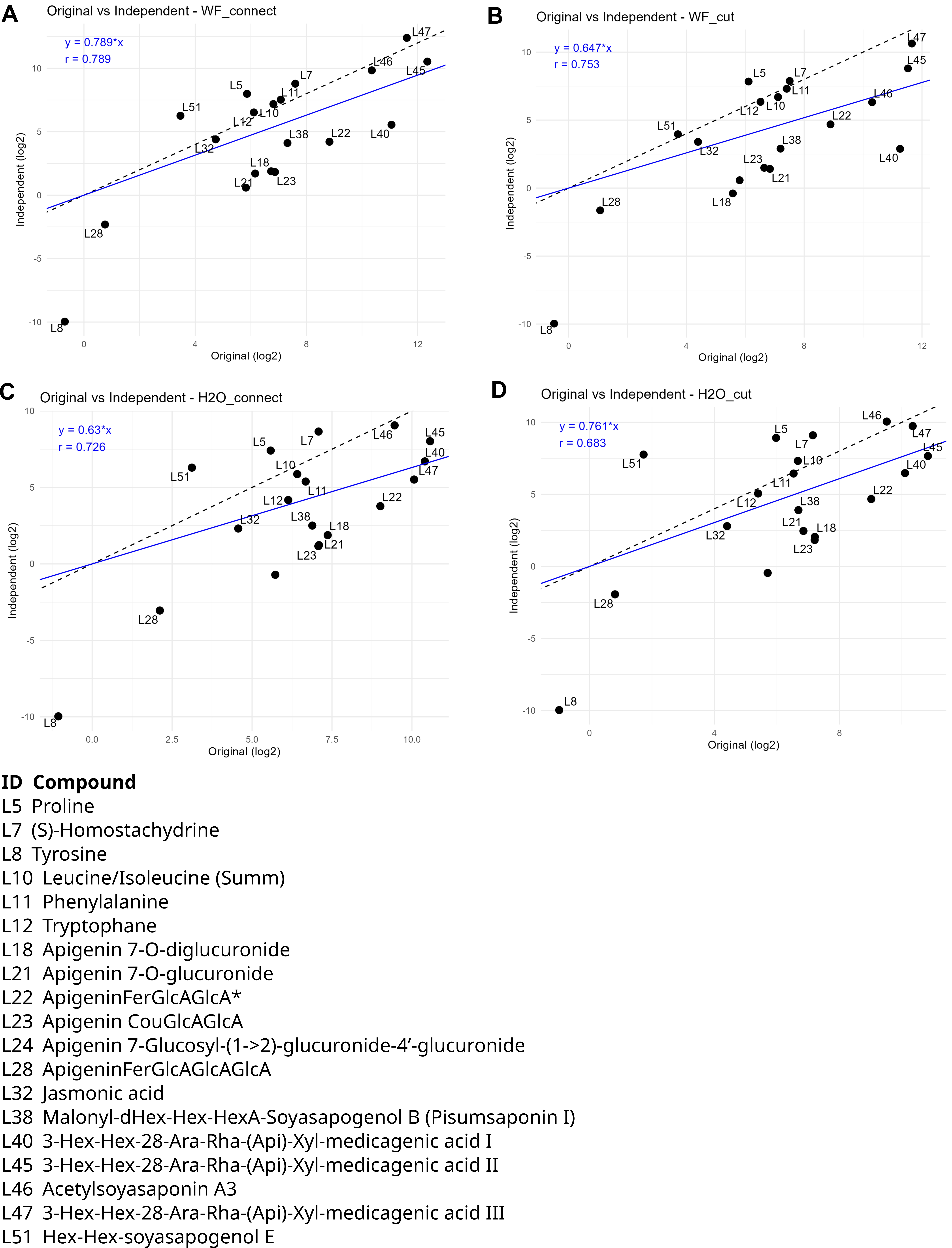
